# NNAlign_MA; MHC peptidome deconvolution for accurate MHC binding motif characterization and improved T cell epitope predictions

**DOI:** 10.1101/550673

**Authors:** Bruno Alvarez, Birkir Reynisson, Carolina Barra, Søren Buus, Nicola Ternette, Tim Connelley, Massimo Andreatta, Morten Nielsen

## Abstract

The set of peptides presented on a cell’s surface by MHC molecules is known as the immunopeptidome. Current mass spectrometry technologies allow for identification of large peptidomes, and studies have proven these data to be a rich source of information for learning the rules of MHC-mediated antigen presentation. Immunopeptidomes are usually poly-specific, containing multiple sequence motifs matching the MHC molecules expressed in the system under investigation. Motif deconvolution -the process of associating each ligand to its presenting MHC molecule(s)-is therefore a critical and challenging step in the analysis of MS-eluted MHC ligand data.

Here, we describe NNAlign_MA, a computational method designed to address this challenge and fully benefit from large, poly-specific data sets of MS-eluted ligands. NNAlign_MA simultaneously performs the tasks of i) clustering peptides into individual specificities; ii) automatic annotation of each cluster to an MHC molecule; and iii) training of a prediction model covering all MHCs present in the training set.

NNAlign_MA was benchmarked on large and diverse datasets, covering class I and class II data. In all cases, the method was demonstrated to outperform state-of-the-art methods, effectively expanding the coverage of alleles for which accurate predictions can be made, resulting in improved identification of both eluted ligands and T cell epitopes. Given its high flexibility and ease of use, we expect NNAlign_MA to serve as an effective tool to increase our understanding of the rules of MHC antigen presentation and guide the development of novel T cell-based therapeutics.

**One Sentence Summary:** Semi-supervised deconvolution and integration of multi-allelic MHC peptidome data allows for improved MHC antigen presentation and T cell epitope predictions.

## Introduction

Major Histocompatibility Complex (MHC) molecules play a central role in the cellular immune system as cell-surface presenters of antigenic peptides to T cell receptors (TCR). Upon presentation, the peptide-MHC complex (pMHC) is scrutinized by T cells and an immune response can be initiated if interactions between the pMHC and TCR are established.

The collection of peptides presented by MHC molecules is referred to as the immunopeptidome. Because of the extreme polymorphism of the MHC, immunopeptidomes can vary dramatically within a population, contributing to the personalized attributes of the vertebrate immune system.

Due to the essential role of the MHC in defining immune responses, large efforts have been dedicated to understanding the rules that shape the immunopeptidome, as well as its alterations in disease – either as a result of pathogen infection or cancerous mutation (*1*). A crucial step towards defining the immunopeptidome of an individual is the characterization of the binding preferences of MHC molecules. The peptide-binding domain of MHC molecules consists of a groove, with specific amino acid preferences at different positions. MHC class I, by and large, loads peptides between eight and thirteen residues long (*2, 3*). MHC class II molecules have an open binding groove at both ends and can bind much longer peptides, and even whole proteins (*4, 5*).

Peptide-MHC binding affinity (BA) assays represented the first attempts of studying binding preferences of different MHC molecules *in* vitro (*6, 7*). However, BA characterization ignores many *in vivo* antigen processing and presentation features, such as protein internalization, protease digestion, peptide transport, peptide trimming, and the role of different chaperones involved in the folding of the pMHC complex (*8*). Further, BA assays most often are conducted one peptide at a time, thus becoming costly, time-consuming, and low-throughput.

Recently, advances in liquid chromatography mass spectrometry (in short, LC-MS/MS) technologies have opened a new chapter in immunopeptidomics. Several thousands of MHC-associated eluted ligands (in short, EL) can with this technique be sequenced in a single experiment (*9*) and numerous assessments, has proven MS EL data to be a rich source of information for both rational identification of T cell epitopes (*10, 11*) and learning the rules of MHC antigen presentation (*12, 13*).

In this context, we have demonstrated how a modeling framework that integrates both BA and EL data achieves superior predictive performance for T cell epitope discovery compared to models trained on either of the two data types alone (*13, 14*). In these studies, the modeling framework was an improved version of the NNAlign method (*15*), which incorporated two output neurons to enable training and prediction on both BA and EL data types. In this setup, weight-sharing allows information to be transferred between the two data types resulting in a boost in predictive power. For MHC class I, we have demonstrated how this framework can be extended to a pan-specific model, capturing the specific antigen presentation rules for any MHC molecule with known protein sequence, including molecules characterized by limited, or even no, binding data (*14, 16, 17*).

Except genetically engineered cells, all nucleated cells express multiple MHC-I alleles and all antigen presenting cells additionally express multiple MHC-II alleles on their surface. The antibodies used to purify peptide-MHC complexes in MS EL experiments are mostly pan- or locus-specific, and the data generated in an MS experiment are thus inherently poly-specific – i.e. they contain ligands matching multiple binding motifs. For instance, in the context of the human immune system, each cell can express up to six different MHC class I molecules, and the immunopeptidome obtained using MS techniques will thus be a mixture of all ligands presented by these MHCs (*12*). The poly-specific nature of MS EL libraries constitutes a challenge in terms of data analysis and interpretation, where, in order to learn specific MHC rules for antigen presentation, one must first associate each ligand to its presenting MHC molecule(s) within the haplotype of the cell line.

Several approaches have been suggested to address this task, including experimental setups that employ cell lines expressing only one specific MHC molecule (*10, 18*–*20*), and approaches inferring MHC associations using prior knowledge of MHC specificities (*21*) or by means of unsupervised sequence clustering (*22*). For instance, GibbsCluster (*23, 24*) has been successfully employed in multiple studies to extract binding motifs from EL datasets of several species, both for MHC class I and MHC class II (*5, 25*–*27*). A similar tool, MixMHCp (*22*) has been applied to the deconvolution of MHC class I EL data with performance comparable to GibbsCluster.

However, neither of these methods are able to fully deconvolute the complete number of MHC specificities present in each dataset, especially for cell lines containing overlapping binding motifs and/or lowly expressed molecules (as in the case of HLA-C). Moreover, for both methods the association of each of the clustered solutions to a specific HLA molecule must be guided by prior knowledge of the MHC binding motifs, for instance by recurring to MHC-peptide binding predictions (*16*). Therefore, both methods require some degree of manual intervention for deconvolution and allele annotation.

A recently published method was suggested to overcome this limitation. The computational framework by Bassani-Sternberg et al. (*28*) employs MixMHCp (*22*) to generate peptide clusters and binding motifs for a panel of poly-specificity MS datasets, and next links each cluster to an HLA molecule based on allele co-occurrence and exclusion principles. While this approach constitutes a substantial step forward for aiding the interpretation of MS EL data, it is clear that it has some substantial shortcomings. First and foremost, the success of the method is tied to the ability of MixMHCp to identify all the binding motifs in a given MS dataset, an ability that is challenged in particular for cell lines containing MHC alleles with similar binding motifs, and for molecules characterized by low expression levels (*22, 29*). Secondly, successful HLA labeling of the obtained clusters is limited by allele co-occurrences and exclusions across multiple cell line datasets. While one may argue that this shortcoming is destined to wane as more immunopeptidomics datasets are accumulated in public databases, there currently remain multiple cases when co-occurrence and exclusion principles fail to completely deconvolute peptidome specificities (*28*).

Inspired by the framework outlined by Bassani-Sternberg et al. *(28*) and by the earlier success of the pan-specific NNAlign framework for modeling peptide-MHC binding (*14*), we here present a novel machine learning algorithm resolving these shortcomings, enabling a fully automated clustering and labeling of MS EL data. The algorithm is an extension of the NNAlign neural network framework (*15, 30, 31*), and is capable of taking a mixed training set composed of single-allele (SA) data (peptides assigned to single MHCs) and multi-allele (MA) data (peptides with multiple options for MHCs assignments) as input and deconvolute the individual MHC restriction of all MA peptides while learning the binding specificities of all the MHCs present in the training set. Compared to earlier approaches for peptidome deconvolution, annotation, and prediction model training (e.g. GibbsCluster/NNAlign (*29*) and MixMHCp/MixMHCpred (*28*)), NNAlign_MA performs these three tasks simultaneously, by iteratively updating the clustering, MHC annotation and peptide binding predictions in an integrated framework. NNAlign_MA does not require manual curation to assign the correct number of clusters, nor for the annotation of clusters to their respective MHC molecule. NNAlign_MA is available at: www.cbs.dtu.dk/suppl/immunology/NNAlign_MA/NNAlign_MA_testsuite.tar.gz.

## Results

A key issue associated with the interpretation and analysis of LC-MS MHC eluted ligand datasets (EL data) stems from the challenge of deconvoluting and linking each ligand back to the presenting MHC molecule(s) of the investigated cell lines. In the following, we describe the NNAlign_MA ramework resolving this challenge, and showcase how the framework can be applied to effectively integrate MA EL data in a semi-supervised manner into machine-learning models for improved prediction of MHC antigen presentation and T cell epitopes on the three large datasets of human (class I and class II) and cattle MHC class I ligand and T cell epitope data.

### The NNAlign_MA algorithm

The NNAlign_MA algorithm is an extension of the NNAlign neural network framework, and is capable of taking a mixed training set composed of single-allele data (SA, peptides assigned to single MHCs) and multi-allele data (MA, peptides that are assigned to multiple MHCs), and fully deconvolute the individual MHC restriction of all MA peptides, learning the binding specificities of the MHCs present in the training set.

The algorithm consists of various critical steps (see Figure 1 for a schematic overview). First, a neural network is pre-trained on SA data only during a burn-in period, using the NNAlign framework underlying the NetMHCpan-4.0 method, which integrates binding affinity (BA) and EL data with sequence information of the MHC molecules. This results in a pan-specific model with potential to infer binding specificities also for MHC molecules not included in the SA dataset (*14, 32*). After this initial training period (from now on referred to as “pre-training”), the data in the MA datasets are annotated. That is, binding for each positive peptide in the MA dataset is predicted (using the ligand likelihood prediction value from the pre-trained model) to all the possible MHC molecules of the given cell line and the restriction is inferred from the highest prediction value (for details see Materials and Methods). For negative MA data, a random MHC molecule from the given cell line is tagged. Next, the SA and now single-MHC annotated MA data are merged, and the model is retrained on the combined data. Note, that the MHC allele annotation is updated at each iteration, and will in general change as the training progresses. Implicitly, the algorithm exploits the principles of co-occurrence and exclusions outlined by Bassani-Sternberg et al. (*28*): i.e. sequence motifs that consistently occur across multiple cell lines sharing only specific MHC alleles are assigned to the shared MHC(s) by the iterative annotation step. For an illustration of this effect refer for instance to HLA-B*13:02, the only allele in common between the two cell lines CM467 and pat-N52. The binding motif for this molecule (and the other HLA molecules in each cell line) as obtained by NNAlign_MA are shown in Supplementary Figure 1. Here, it is apparent that only one motif is shared by these two cell lines, and the co-occurrence principle allows NNAlign_MA to assign this motif to HLA-B*13:02. In ambiguous cases where co-occurrence and exclusion principles are insufficient, the pan-specific nature of the method will help tilt the annotation towards the correct MHC by similarity to already characterized, unambiguous MHC alleles. An example of this is the BoLA-1:00901 case in Figure 1, where H at P9 - even though only marginally tolerated after pre-training - fits better to this molecule compared to the alternative allelic options in the relevant cell lines, thus tilting the ligands with H at P9 towards this molecule. For details on model hyper-parameters and model training setup, see Supplementary Material.

**Figure 1.**
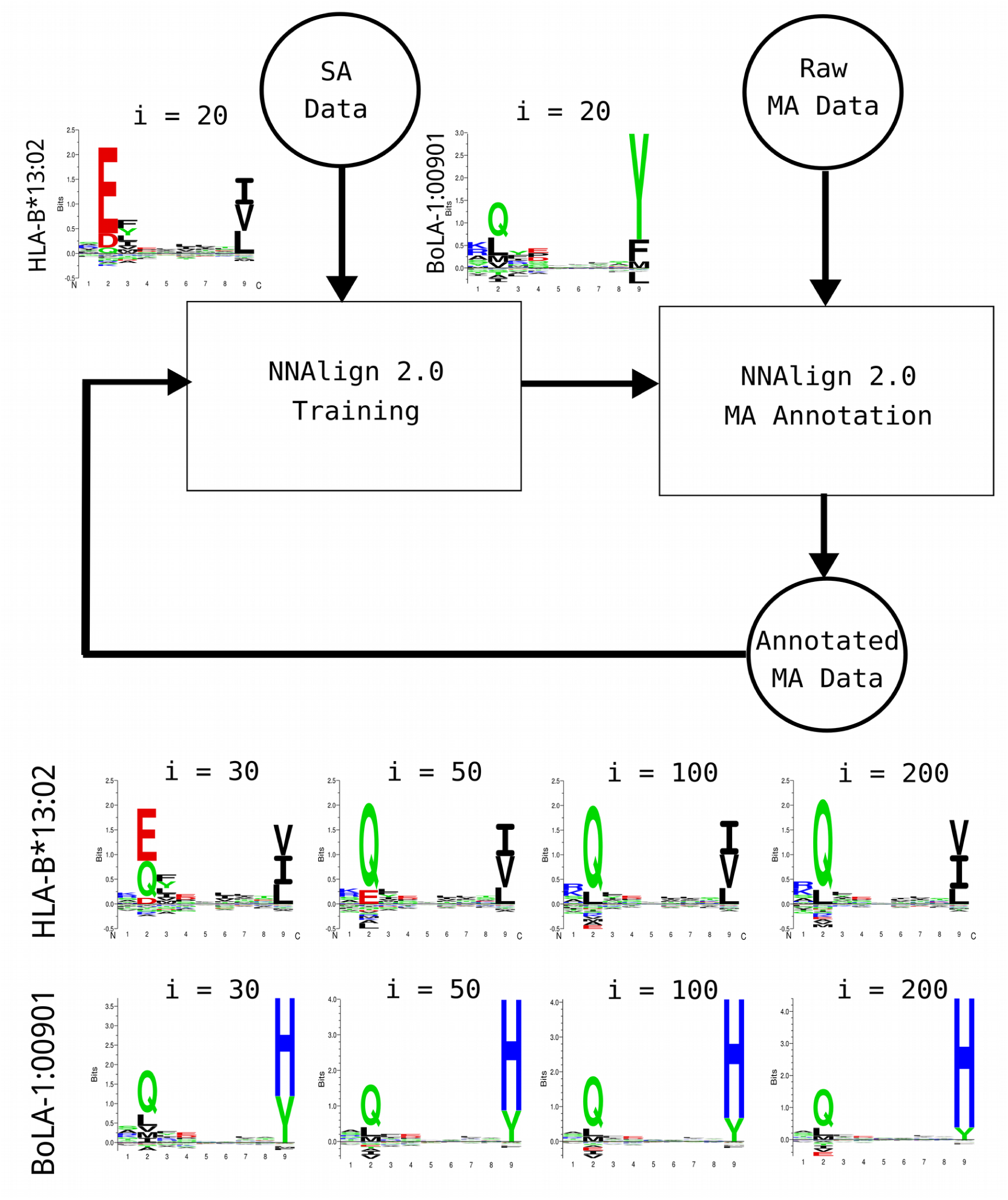
NNAlign_MA framework. An initial model is pre-trained using SA data, next MA data are annotated and merged with the SA data and the training repeated iteratively. In the upper left part are displayed examples of binding motifs of two MHC molecules after pre-training (i=20). In the lower part of the figure are shown the changes to predicted binding motifs of the same two MHC molecules as NNAlign_MA iterates over the data. Here i refers to the number of iterations.

### HLA-I benchmark

To benchmark this novel approach, we trained a model on the complete set of SA data described in the NetMHCpan-4.0 paper, combined with an extensive set of MA data covering 50 different cell lines with typed HLA allotypes described in Bassani-Sternberg et al. (*28*). Note that, prior to training, we removed all overlapping peptides between the SA and MA data from the SA data set. This was done to fully demonstrate the power of the NNAlign_MA to annotate MA data also in situations where the information cannot simply be transferred from the SA data. After training, each MA data point ends up being annotated to a single MHC molecule, and from this annotation the distribution of ligands associated with each HLA molecules was recovered. Based on the predicted 9mer binding cores of the ligands, logos for all the HLA alleles expressed by each of the cell lines under study was constructed (Supplementary Figure 1). In this benchmark, NNAlign_MA was capable of not only clustering the EL data into a set of groups matching the number of expressed HLA alleles in each cell line (this is guaranteed by the construction of the method), but also to assign each group to a single corresponding HLA allele. As a point of comparison, on the same benchmark data, MixMHCp was only capable of achieving a complete deconvolution of all HLA specificities in 26% of the 50 cell line data sets (failing to identify motif corresponding to HLA-C alleles in 61% of the cases), and could not annotate at least one cluster in 16% of the samples. Two examples of this are given in Figure 2, showing the NNAlign_MA deconvolution of the two cell lines HCC1143 and HCT116. In the first case, MixMHCp correctly identified 5 motifs, but could not assign two of the five to their corresponding allele (respectively HLA-A*31:01 and HLA-B*37:01). For HCT116, MixMHCp was able to deconvolute and assign only four of the six expressed alleles, missing the deconvolution of the motifs for two HLA-C alleles, HLA-C*05:01 and HLA-C*07:01. The accuracy of the 4 motifs additionally identified by NNAlign_MA was confirmed by reference to SA data available from IEDB (*33*) (see Figure 2).

**Figure 2.**
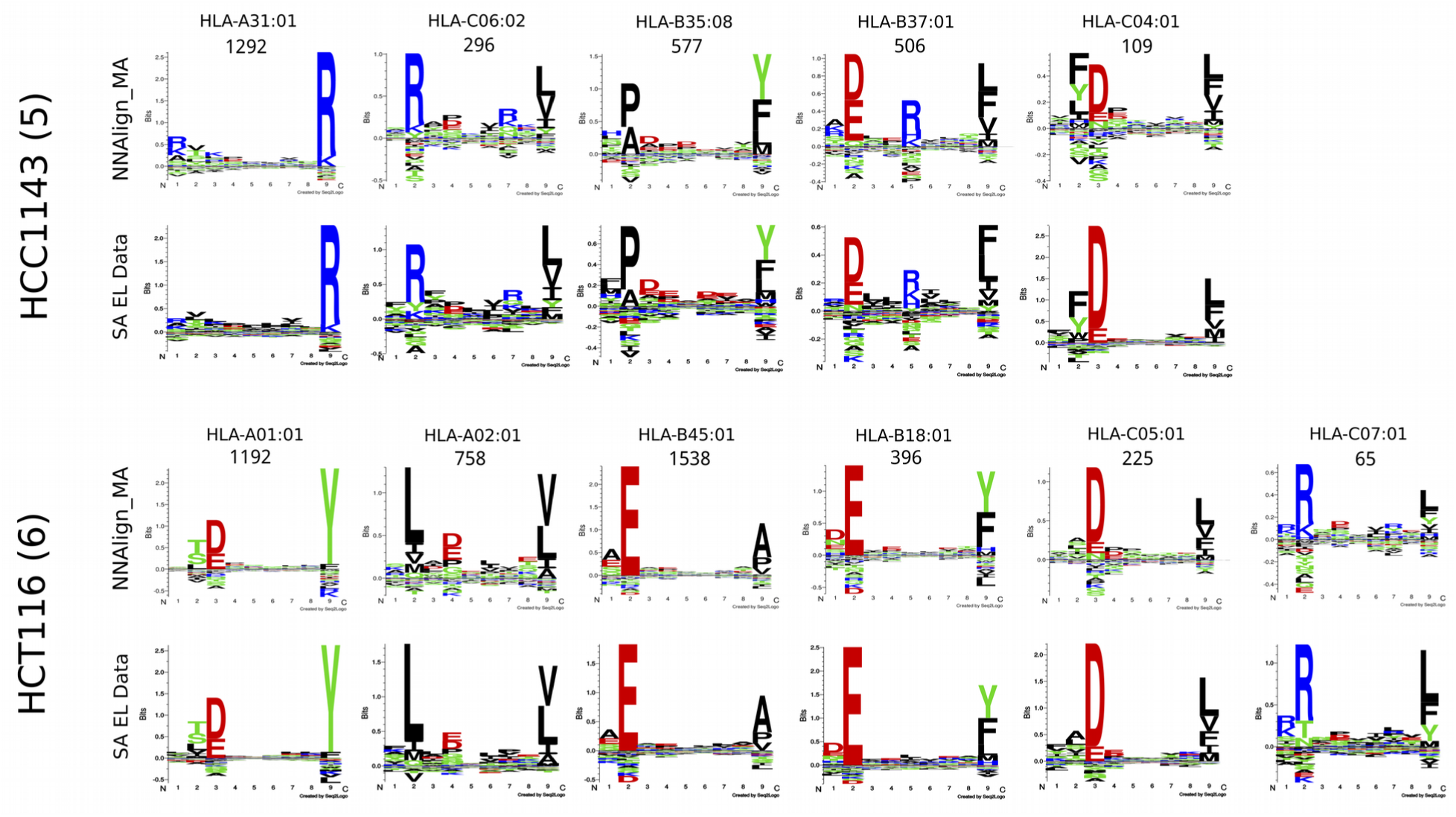
Sequence clustering and labeling comparison between NNAlign_MA and NetMHCpan-4.0 for the HCC1143 and HCT116 cell lines. Motifs corresponding to NNAlign_MA were constructed based on ligands from the given MA data set (cell line) predicted using cross-validation to be restricted by the given HLA molecule; the quantity of peptides associated to each HLA molecule is given on top of the corresponding logos. SA EL data motifs were derived from single-allele (SA) data available from the IEDB (*33*).

As stated above, the NNAlign_MA method by construction is guaranteed to cluster the MA data into a number of groups matching the number of HLA alleles expressed in each cell line. The association of each cluster to the correct HLA molecule, and the accuracy of each cluster are, however, not guaranteed. By investigating the deconvolution solutions for the different cell lines (Supplementary Figure 1), it is apparent that the accuracy of the motifs identified by NNAlign_MA (as expected) depends on the number of ligands assigned to a given HLA, and that complete characterization of the HLA’s in a given cell line, for a few cases, is impeded by this fact (a few examples include the motif for HLA-C*07:04 from Fibroblast, HLA-C*08:01 from Mel-624, and HLA-C*02:10 from RPMI8226). These are further examples of alleles only present in single MA data sets, limiting the ability of NNAlign_MA to transfer information of the binding motifs from other data sets.

To further quantify the accuracy of the cluster-HLA association, we compared the motifs obtained by NNAlign_MA to the motifs obtained from SA data in situations where such data were available from the IEDB (Supplementary Figure 2). Here, we in the vast majority of cases observed an excellent agreement, with an average correlation between the two motifs of 0.869 over the 49 alleles included (p-value for the correlation being random was in each case < 0.001, exact permutation test, for details on how the correlation was calculated refer to Supplementary Materials). Note, that this correlation was equally high for alleles belonging to the HLA-A/B, and HLA-C loci. As expected, the agreement between the MA and SA motifs also here was highest for the cases where both motifs were characterized by large data sets.

Next, we compared the motifs of individual HLA alleles obtained across different cell lines, for example the HLA-C*03:03 allele sharece when evaluated on the SA data (median AUC of 0.9842 versus 0.9839). We hypothesized d between 5 cell lines. Using again a simple correlation analysis, we quantified the similarity of these different motifs, and could in all cases confirm a high consistency, with an overall averaged correlation of 0.901 over the 17 alleles shared by 5 of more cell lines (Supplementary Figure 3). These correlation values were all significantly different from random (p<0.0001, exact permutation test), and significantly higher than the correlations obtained by comparing motifs assigned to different HLA molecules (p<10^-5^, T-test). Also, the correlations were lowest for the comparison between motifs characterized by small data sets (as exemplified by the motifs for HLA-C*07:02 from the HCC1937 and Mel-8 cell lines, each characterized by 48 and 31 ligand data points respectively (see Supplementary Figure 1), resulting in a correlation between the two motifs of 0.68). Finally, we evaluated the “cleanness” of each cluster/motif by calculating predicted positive (PPV) values. Here, all clustering solutions were found to have very high accuracy, with an average PPV value of 75% (for details on the calculation of the PPV refer to Supplementary Material, and for the complete list of PPV values refer to Supplementary Table 1).

Taken together, these results demonstrate the high performance of NNAlign_MA, achieving in the vast majority of cases an accurate, consistent and complete (including for HLA-C) deconvolution of MA EL data sets.

Given these encouraging results, we next conducted a full-scale performance evaluation for prediction of eluted HLA ligands. To this end, we first compared the performance of NNAlign_MA trained on the complete HLA-I dataset (referred to as the MA model) to the performance when trained only on the subset of SA data (referred to as the SA model). This benchmark (Figure 3A) demonstrated that the MA model exhibited a consistently (and statistically significant, p<0.0001, paired T test) higher performance when evaluated on the MA data (median AUC of 0.9769), in comparison to the SA model (median AUC of 0.9712). On the other hand, as expected, the MA and SA models showed an overall comparable predictive performanthat NNAlign_MA would demonstrate a performance gain over NNAlign_SA for alleles where the SA data are either limited or absent. The results display in Figure 3 confirmed this. Here, the median number of positives for SA data sets where NNAlign_MA outperforms NNAlign_SA was 57 whereas the number for the SA data sets where NNAlign_SA won was 435. Further, was the performance gain of NNAlign_MA on the MA data found to be largest for data sets characterized by alleles absent from the SA data.

**Figure 3.**
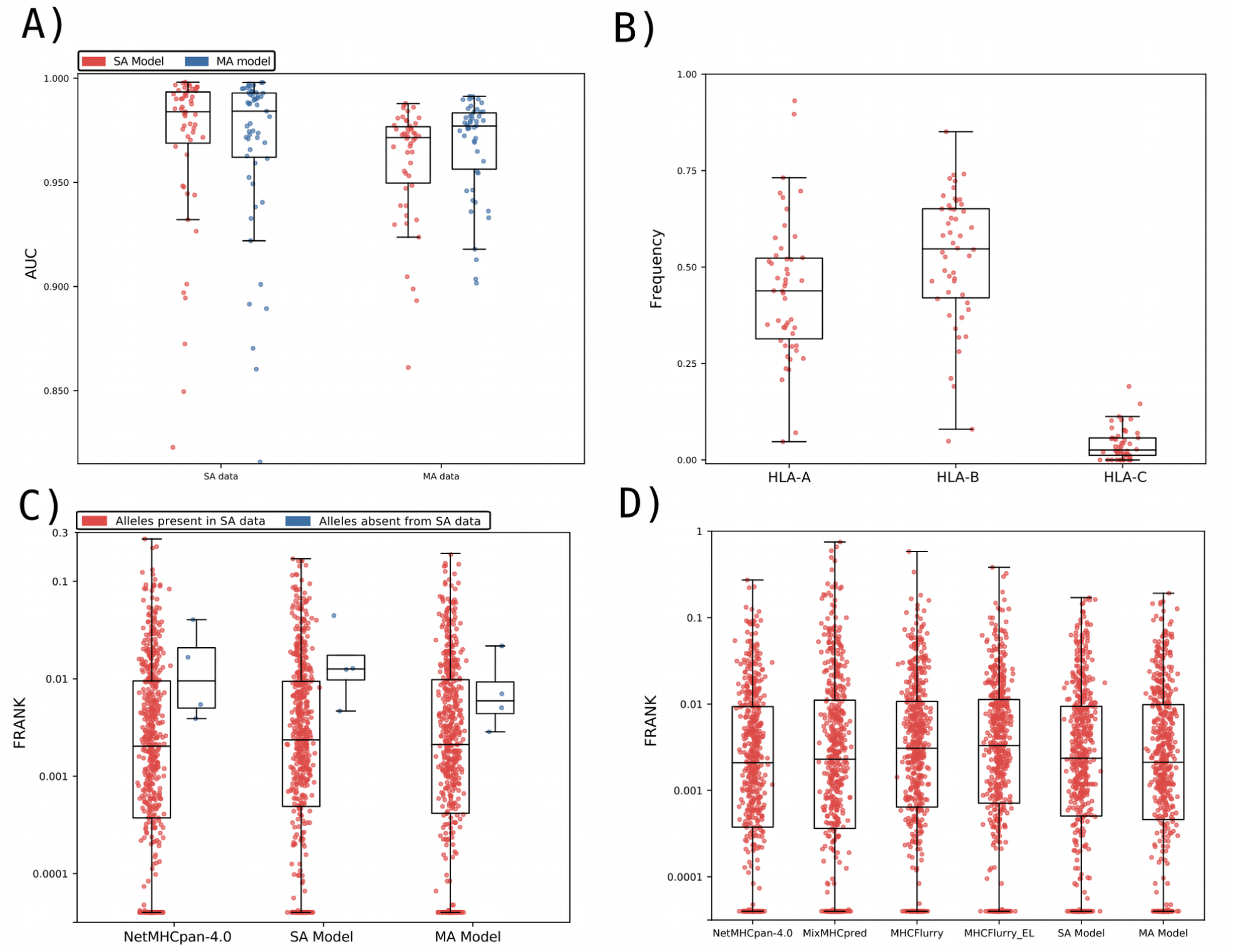
Benchmark of the prediction method on the HLA-I data. A) Performance of the SA and MA trained models on the SA EL and MA EL data sets, expressed in terms of AUC. Each point corresponds to one SA or MA data set. SA data corresponds to 55 single-allele EL dataset. MA data consist of EL data from 50 different cell lines, each expressing more than one MHC molecule. To evaluate the MA data, each data point was assigned the highest prediction value across all possible MHC restrictions in the given cell line dataset (for further details, see Materials and Methods). Performance values for the SA model on the SA data sets, and for the MA model on the SA and MA data, were extracted from the cross-validated predictive performance. B) The relative peptidome size for the three loci (HLA-A, HLA-B and HLA-C) as predicted by NNAlign_MA for the different MA datasets. Each data point gives the relative proportion of ligands in each MA dataset predicted to be restricted by a HLA from the given locus. The HLA restriction for each ligand data was estimated from the evaluation performance of the cross-validation as described in Material and Methods. Only datasets where HLA expression is annotated for all three loci were included. C) Frank values for the epitope evaluation dataset for NetMHCpan-4.0 and models trained with only SA data (SA Model) and with SA and MA data (MA Model). Red dots correspond to epitopes restricted to HLAs that were part of the SA training data, while blue points refer to Frank values for epitopes with HLA restrictions absent from the SA training set. For visualization, Frank values of 0 are displayed with a value of 0.00004. HLAs in the category “Alleles absent from SA data” are HLA-B*13:02, HLA-B*55:01 and HLA-C*01:02. D) Frank values for the epitope evaluation on NetMHCpan-4.0, MixMHCpred, MHCFlurry (trained on BA data), MHCFlurry_EL (trained on BA and EL data) and the models trained with only SA data (SA Model) and MA data (MA Model). For visualization, Frank values of 0 are displayed with a value of 0.00004.

Next, we investigated how the peptidome for the MA EL data in each dataset was distributed among the alleles of the three loci HLA-A, HLA-B and HLA-C. To do this, we extracted the number of ligands predicted by NNAlign_MA to be restricted by HLA-A, HLA-B and HLA-C for each cell line present in the MA EL dataset and then calculated the proportion of ligands associated to a given loci relative to the total amount of peptides in the cell line. The result of this analysis is shown in Figure 3B, and confirms the general notion that HLA-A and HLA-B have comparable peptidome repertoire size, whereas the peptidome size of HLA-C, in comparison, is substantially reduced (*34*). While this is not a novel observation, to the best of our knowledge this is the first fully automated analysis of EL data demonstrating this. Some clear outliers are present in the figure where either the HLA-A or HLA-B peptidome repertoires are highly reduced compared to the median values. A few such examples include the HL-60, CA46, Mel-624 and HEK293 cell lines where either the entire HLA-A or HLA-B locus appears to have been deleted or made non-functional. These observations are in agreement with results from earlier studies for these cell lines (for details refer to Supplementary Table 2), suggesting the power of NNAlign_MA also for identification of loss or down-regulation of HLA expression directly from EL datasets.

### Evaluation on HLA-I epitopes from IEDB

To further investigate the predictive power of NNAlign_MA, we employed an evaluation set of epitopes of length 8-14 extracted from IEDB (see Supplementary Material for details). Here, we divided the dataset into two subsets; one containing the epitopes restricted to HLAs that were part of the SA training dataset, and one where the HLAs were not present in the SA training data. The results of the evaluation on these two datasets for the SA and MA models, and the state-of-the-art method NetMHCpan-4.0 are shown in Figure 3C in terms of Frank values. This measure reflects the false-positive rate, and a value of 0 corresponds to the perfect prediction (for details see Supplementary Material). For epitopes restricted to HLA molecules that are part of the SA dataset (red points), the figure displays a comparable performance of the MA and NetMHCpan-4.0 methods (median Frank values 0.0021 and 0.0020, p=0.3, paired T test), and a significantly worse performance of the SA method (median Frank 0.00218, p<0.0025 paired T test). Further, when evaluated on the small set of epitopes whose HLA restrictions are only present in the MA data (blue points), the performance of the MA model is substantially increased compared to both the SA and NetMHCpan-4.0 models (average Frank of 0.0091 compared to 0.0186 and 0.0166 for the SA and NetMHCpan models). These results demonstrate how the NNAlign_MA model also when it comes to prediction of T cell epitopes achieved state-of-the-art performance, and further is capable of benefiting from MA data to expand the allelic coverage outside SA data set to improve the allelic coverage and predictive power.

Finally, in figure 3D, the evaluation was expanded to include the MHCFlurry (trained without and with EL data) and MixMHCpred methods limiting the benchmark to include only HLA molecules covered by all methods, thus including only HLA alleles with previously well-characterized binding motifs (for details on the benchmark refer to Supplementary Material). The results of this benchmark confirmed a comparable performance of NNAlign_MA (median Frank 0.0021) to NetMHCpan-4.0 (median Frank 0.0021), and a small (but statistically significant, p<0.05 paired T-test, in all cases except for MHCFlurry) drop in performance of MHCFlurry (median Frank 0.0031), MHCFlurry_EL (median Frank 0.0033), MixMHCpred (median Frank 0.0023) and NNAlign_SA (median Frank 0.0024). This result confirms the state-of-the-art performance of NNAlign_MA.

### A specificity leave-out benchmark

All the benchmarks performed hereto were conducted in situations where the MA data shared high HLA overlap with the SA data. By way of example, over 75% (51 of 67) of the alleles in the MA dataset were part of the SA data, and 94% (63 of 67) share a distance of less than 0.1 to an allele in the SA data as measured from the similarity between pseudo sequences (for details on this similarity measure see Supplementary Material), a distance threshold earlier demonstrated to be associated with high predictive accuracy of the pan-specific prediction model (*35*).

As stated above and confirmed by the results in Fig 3A and 3C, the main power of NNAlign_MA is to effectively extend the allele-space covered by HLA annotated EL data leading to an improved predictive power outside the space covered by SA data. Given the high allelic overlap between the MA and SA data set, it is hence not surprising that the impact of including MA data in these benchmarks was limited. Therefore, to further test the power of the NNAlign_MA method in a more extreme setting, we conducted an experiment where parts of the SA data were left out from the training data leaving part of the HLA space covered only by MA data. In short, we removed all SA data for HLA molecules belonging to (or similar to alleles in) the HLA-A2 and HLA-A3 supertypes (*36*), effectively pruning off whole branches from the tree of HLA specificities (Figure 4 left panel) (for details on this pruning refer to Supplementary Material). This scenario thus simulates a situation where the MA data describes binding specificities that a novel compared to any specificity contained in SA training data. Given this, the binding motifs in the MA data cannot simply be inferred from a close neighbor in the SA data, making the challenge of HLA deconvolution non-trivial. This experiment therefore allows us to investigate how the NNAlign_MA method can benefit from MA data to accurately characterize the binding specificity of HLA molecule not characterized by SA data, and from such MA data expand the HLA coverage of the trained prediction model.

**Figure 4.**
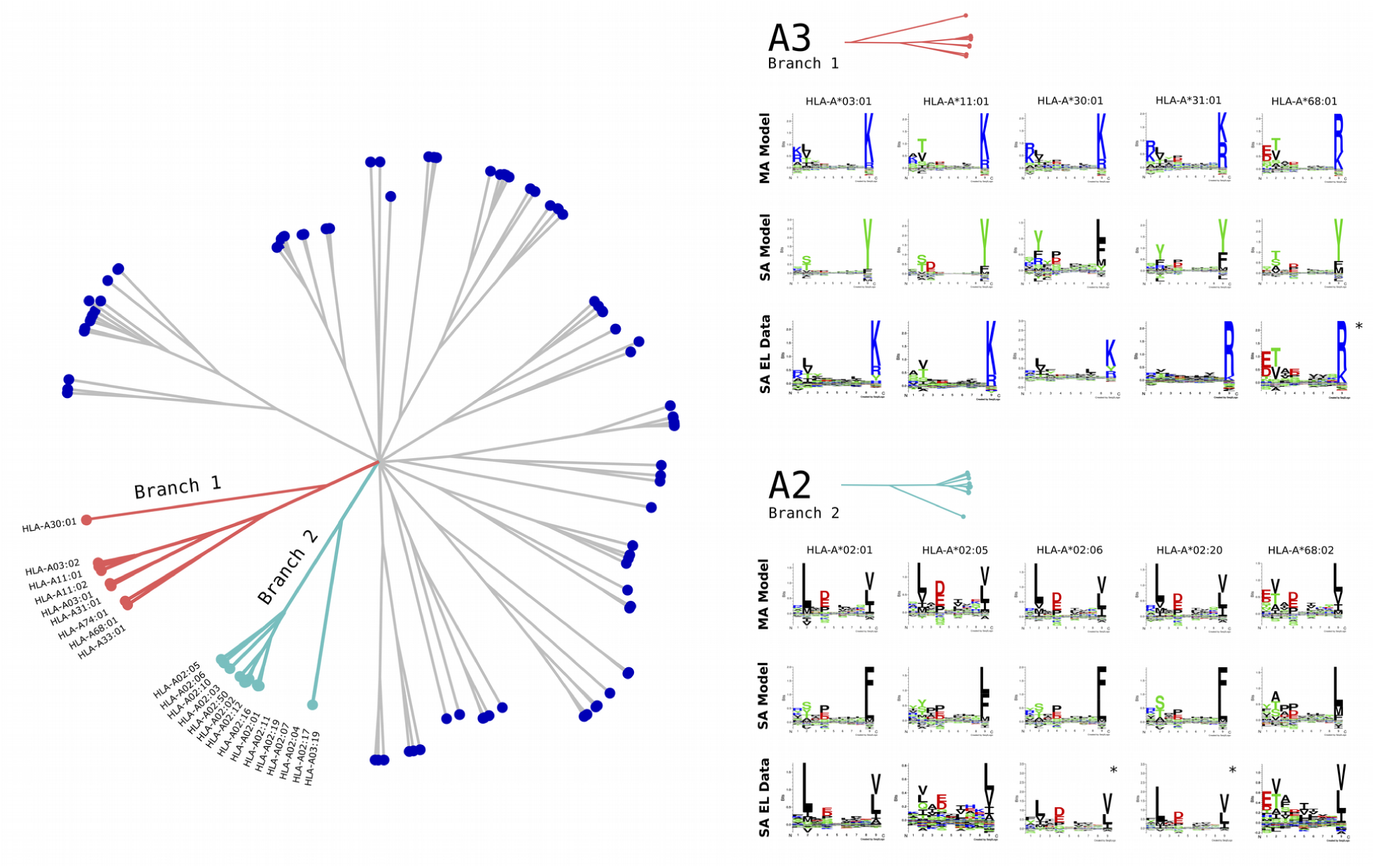
HLA supertype pruning experiment. The left panel shows a functional tree of HLA specificities estimated using MHCcluster (*37*). Branches in light blue and red correspond to the A2 and A3 HLA supertypes. In the HLA supertype tree pruning experiment, SA data for HLA molecules belonging to both these branches were removed. Right panel shows binding motifs for A2 and A3 supertype alleles from the MA EL dataset predicted by the MA and SA models. Motifs were constructed from the top 1% of 1,000,000 random natural 9mer peptides predicted by each model. SA EL data show motifs derived from SA EL data (not included in the training). For alleles marked with * no SA EL data was available and motifs were obtained from NetMHCpan from http://www.cbs.dtu.dk/services/NetMHCpan/logos_ps.php.

In the benchmark, SA and MA models were trained as described above on the pruned SA and complete MA data, and the predictive performance was evaluated on SA EL data for the alleles on the pruned branch. Note, that this evaluation was done respecting the data partitioning of the cross validation to avoid introducing a bias in favor of the MA model. The performance was estimated in terms of AUC0.1 resulting in average values of 0.599 versus 0.852 for the SA and MA models respectively (for details on the performance values see Supplementary Table 3).

Next, binding motifs for the alleles in the MA data from the A2 and A3 supertypes were estimated for the MA and SA models and compared to motifs derived from SA EL data if available (see Figure 4 right panel). Here, we observed a high overlap between the motifs of the MA model motifs obtained from SA EL data, and a likewise low overlap of the motifs obtained by the SA model. Note, also here that the agreement between the MA model and SA EL data motifs was dependent on the number of ligands assigned to the given allele from the MA data, i.e. HLA-A*02:01 was assigned 26,038 and HLA-A*02:05 only 917 ligands from the MA data resulting in a somewhat lower agreement between the two motifs for HLA-A*02:05 compared for HLA-A*02:01. Finally, we reinvestigated the performance of the SA and MA models trained on the pruned SA data set, on the subset of 358 epitopes restricted by the 11 alleles covered by the two A2 and A3 supertypes. This benchmark confirmed the superior performance of the MA model over the SA model with median Frank values of 0.0044 compared to 0.0393 (p<10^-15^, paired T test). Note, the performance of the full MA model on this epitope data set was 0.0032. Taken together these results demonstrate the power of the NNAlign_MA to accurately characterize individual binding motifs of molecules from MA data only.

### BoLA-I benchmark

Having demonstrated how NNAlign_MA was capable of benefitting from MA EL data to boost predictive power and expand the allelic coverage also in a setting where the MA data shared limited allelic overlap to the SA data, we next turned to the BoLA (Bovine Leukocyte Antigen) system. Because binding data (both BA and EL) is more scarce for BoLA compared to HLA, and because the relative expression of MHC molecules within a given cell line varies in a more dramatic manner for the bovine system compared to humans, analyzing and deconvoluting BoLA MA EL data is more challenging compared to HLA, and working within this system allowed us to better appreciate and assess the strength and potential limitations of the NNAlign_MA framework.

In a previous work, a prediction model for BoLA peptide interactions, NetBoLApan, was trained using NNAlign on SA BA (including binding affinity data for 7 BoLA molecules) and EL MA data from 3 BoLA-I homozygous cell lines, describing the BoLA haplotypes A10, A14, and A18 (*16*). Due to the prior limitation of the NNAlign framework only admitting SA data for training, in NetBoLApan the EL MA data had to be first deconvoluted using GibbsCluster and then manually annotated to the individual BoLA molecules of each cell line by visual inspection. Since this earlier publication, we have generated MA EL data for additional 5 cell lines. Using these data, we trained and evaluated the NNAlign_MA framework on the SA data described above combined with EL MA data for a total of 8 BoLA cell lines (for details on these datasets, refer to Supplementary Material).

The benchmark on the BoLA EL data confirmed the power of the NNAlign_MA method to achieve complete, consistent and well-defined deconvolution and motif identification of MHC alleles in the MA EL data sets, also in this challenging case with limited overlap between the SA and MA data (details on this benchmark are found in Supplementary Material, and Supplementary Figure 4). One notable example demonstrating this is BoLA-1:00901, a molecule contained within the A15 haplotype. BoLA-1:00901 shares limited overlap with the SA training data (distance to the SA data D=0.137), and the motif predicted by the NNAlign_MA method after the pre-training on the SA data share, as expected, share high similarity to the motif predicted by NetBoLApan (Figure 5). This pre-trained motif is, however, altered substantially after the training of the model on the EL MA BoLA data, resulting in a strong preference for Histidine (H) at PΩ (Post-training motif in Figure 5). To validate the accuracy of the motif predicted for BoLA-1:00901, we performed *in vitro* binding assays of a combinatorial peptide library to the BoLA-1:00901 molecule (for details see Supplementary Material). The *in vitro* binding motif showed very high similarity to the Post-training motif predicted by NNAlign_MA (Figure 5).

**Figure 5.**
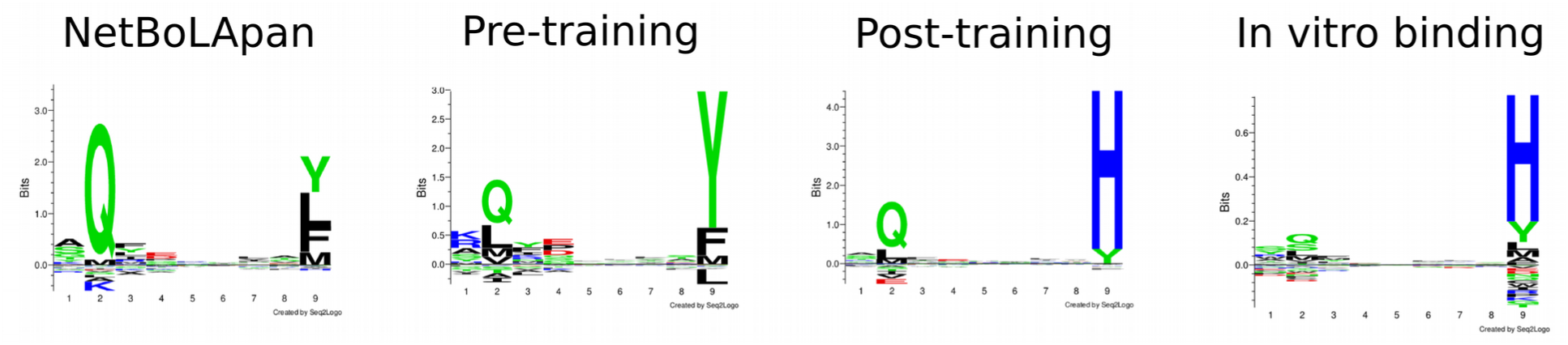
Binding motifs for the BoLA-1*00901 molecule estimated by different *in silico* and *in vitro* binding methods. Binding motifs for the three *in silico* methods were estimated from the top 0.1% of 1,000,000 random natural 9mer peptides with predicted binding by the given method, for BoLA-1:00901. The *in vitro* binding motifs were estimated using a position scanning combinatorial peptide library, as described in Materials and Methods. The three *in silico* methods are: NetBoLApan (*16*), trained including EL data for the cell lines A10, A14 and A18; Pre-training, the NNAlign_MA method pre-trained on SA data; Post-training, the NNAlign_MA method after completing the training including MA data.

### Evaluation on BoLA-I epitopes

Having demonstrated the power of the proposed model also for the challenging BoLA system, we next evaluated its predictive power on a set of experimentally validated BoLA restricted CD8 epitopes. The result of this evaluation for NNAlign_MA and NetBoLApan confirmed the high performance of the proposed model (Table 1). Overall, the performance of the NNAlign_MA model is comparable to that of NetBoLApan. However, one notable example where the two models showed very different performance is the FVEGEAASH epitope, restricted by BoLA-1:00901. For this epitope, the Frank performance value of NetBoLApan was 0.121 - in other words, the true positive is found 12.1% down the list of candidate peptides predicted by NetBoLApan. Including the novel BoLA EL data and training the model using the NNAlign_MA framework, the Frank value for this epitope improved to 0.000 - the epitope is the single top candidate predicted by NNAlign_MA. This result aligns with the experimental binding motif analysis of the BoLA-1:00901 molecule, exhibiting a strong preference for H at the C terminal (Figure 5). In summary, the results displayed in Table 1 demonstrate the high predictive power of the model trained including the BoLA EL data also for prediction of CD8 epitopes. The average Frank value of NNAlign_MA over the 16 epitopes is 0.0033, meaning that on average 99.67% of the irrelevant peptide space can be excluded by the prediction model while still identifying 100% of the epitopes.

**Table 1.**
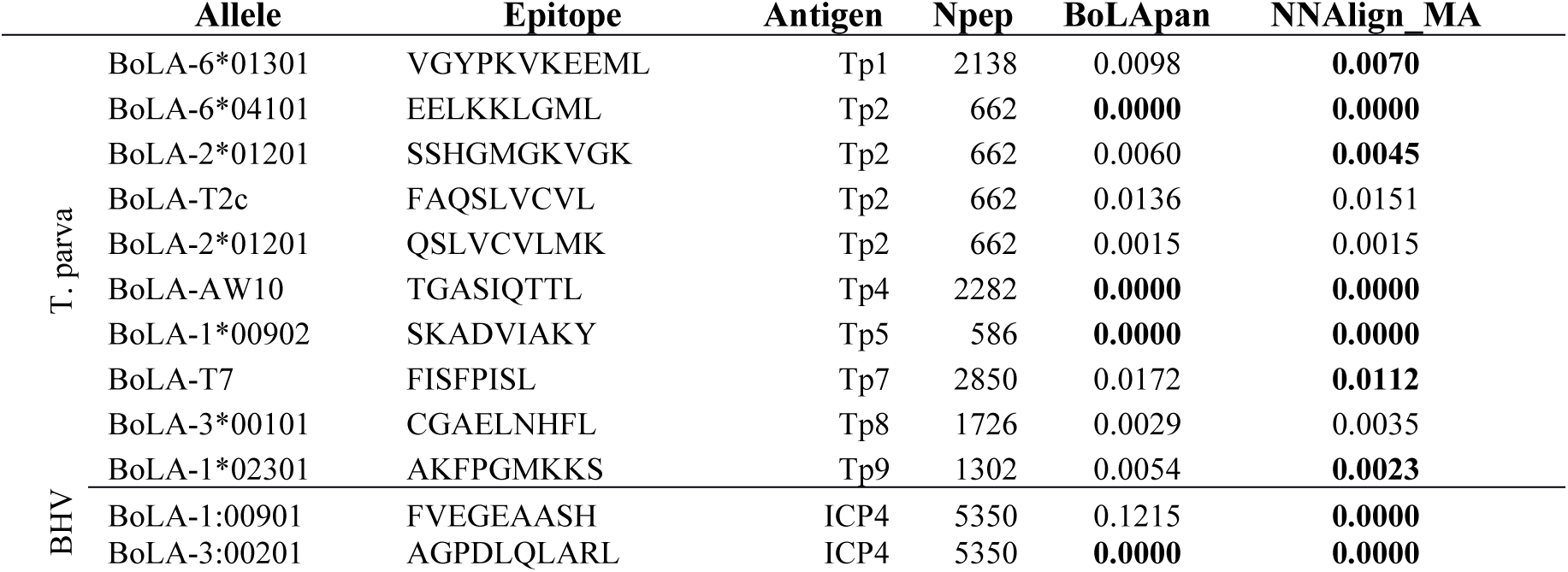
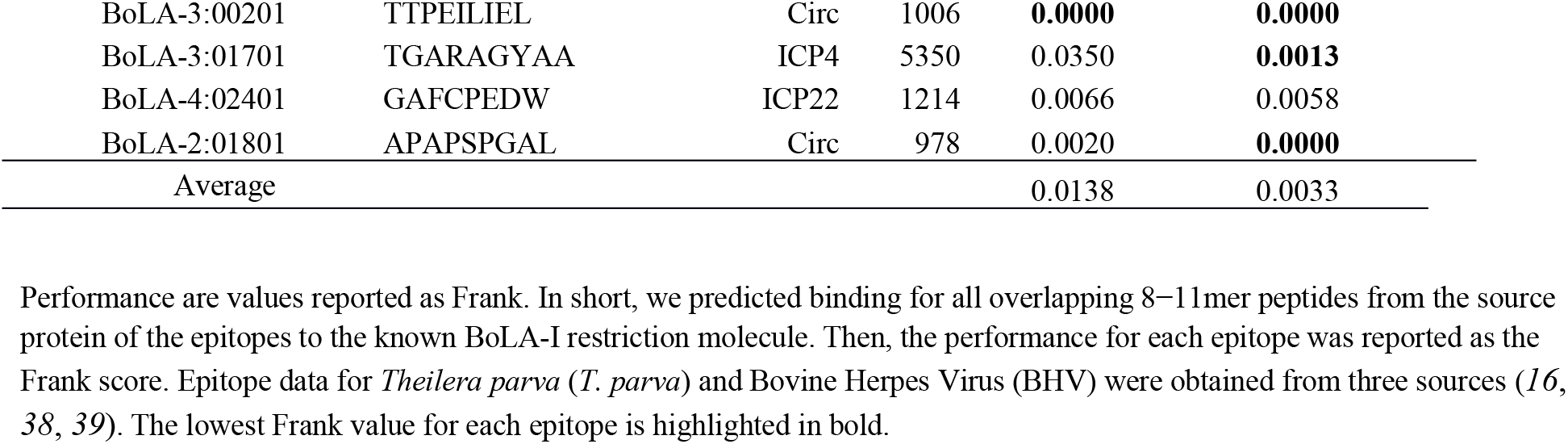
Predictive performance of NNAlign_MA and NetBoLApan on the set of known CD8 epitopes.

### HLA-II benchmark

To prove that the ability of NNAlign_MA to deal with MA data also extended to MHC II, a separate study was conducted on a set of MHC II BA and EL data (for details see Supplementary Material). Here, we compared the cross-validated performance of NNAlign_MA trained on SA data alone (SA model) versus NNAlign_MA trained on the full dataset including MA data (MA model). Both models were evaluated individually on the SA and MA data sets as described earlier for the HLA-I benchmark. The conclusions from this evaluation (Figure 6A) were similar to those obtained for the HLA-I data: when evaluated on SA data, the SA model demonstrated a modest (and statistically insignificant (p=0.125, binomial test) performance gain compared to the MA model. However, when it comes to the MA data, the MA model significantly outperformed the SA model (p=7.6 * 10^-6^, binomial test excluding ties). In Supplementary Figure 5, we further show the binding motifs for MA data included in this study demonstrating that also for class II is the NNAlign_MA framework in the vast majority of cases capable of achieving clear and consistent MHC motif deconvolution. However, as for the class I data does the accuracy of the motifs identified by NNAlign_MA also here depend on the number of ligands assigned to a given HLA. For example, is the motif for HLA-DRB1*13:01 most often derived from a very small number of ligands resulting in a limited similarity between the motifs obtained from the different MA data sets. This observation underlining the critical dependency of NNAlign_MA on the quality of the input data.

**Figure 6.**
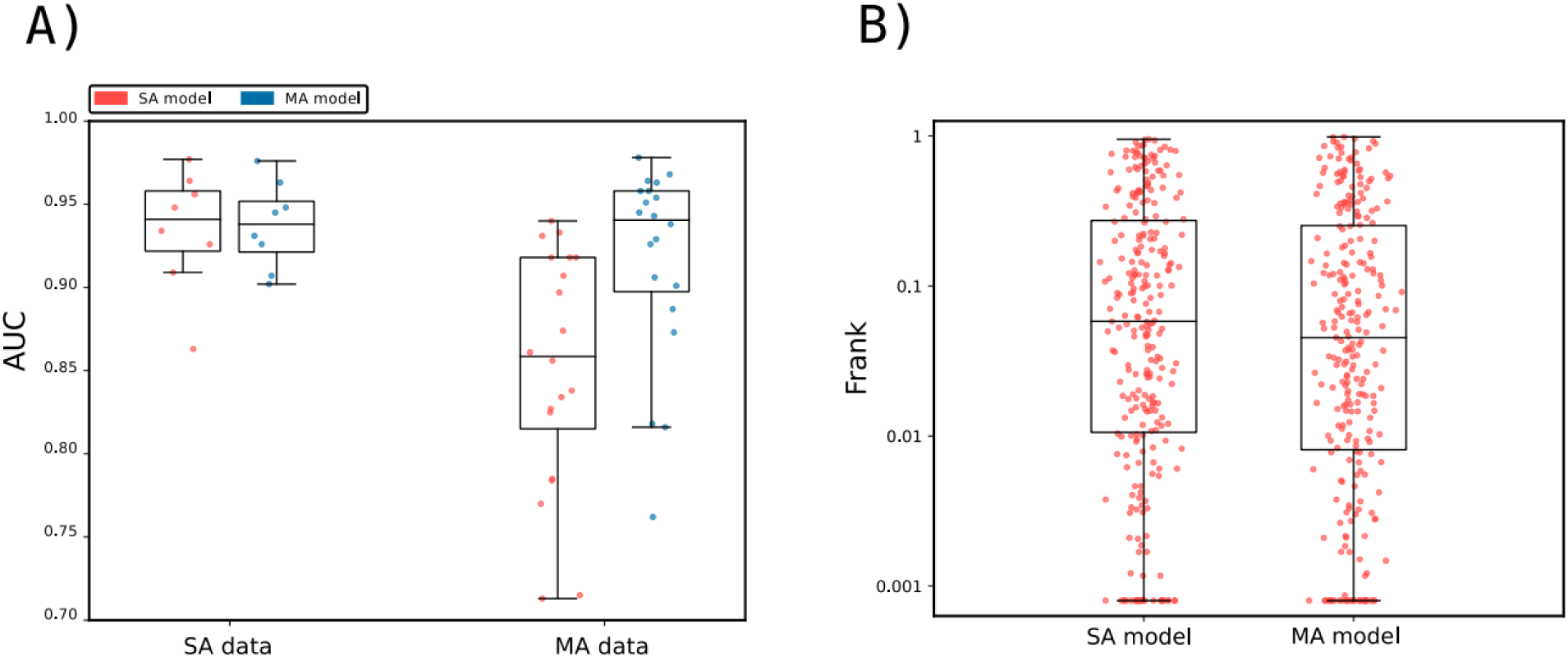
Method benchmarking on HLA class II data. A) Performance of NNAlign_MA trained on SA and MA data evaluated in cross-validation. Model performance is given in terms of AUC and calculated as described in Figure 3 Each point refers to an individual SA or MA data set. B) Epitope Frank scores of NetMHCIIpan-3.2 and NNAlign_MA trained on SA and MA data evaluated on CD4+ epitopes from the IEDB (*33*). The plot compares the Frank distribution of the MA and SA models and each point represents the Frank of an epitope. For visualization, Frank values of 0 are displayed with a value of 0.0008.

Next, we evaluated the performance of the SA and MA models together with NetMHCIIpan-3.2 on an independent data set of CD4+ epitopes (for details on this data set refer to Supplementary Material). The results of this benchmark are depicted in Figure 6B in terms of Frank values. Here, the median Frank values were respectively 0.02403, 0.03155, and 0.03734 for the three methods MA, SA and NetMHCIIpan-3.2. The difference in Frank between the MA and the two other methods was in both cases found to be statistically significant (p<0.01, binomial test excluding ties). These results strongly suggest that the NNAlign_MA framework extends its predictive power also into MHC class II.

## Discussion

Advances in Mass Spectrometry have dramatically increased the throughput of immunopeptidomics experiments, with several thousands of peptides directly eluted from their cognate MHC molecule in a single experiment. This type of data has greatly changed our knowledge base for characterizing MHC antigen processing and presentation. In general, MS eluted ligands originate from multiple MHC molecules, and MS datasets therefore consist of a mixture of motifs, each corresponding to the binding specificity of one of the MHC molecules expressed by the cell line. While several tools for the deconvolution of multiple motifs have been proposed, they all tend to underestimate the number of specificities in a sample, especially for haplotypes with overlapping MHC binding motifs and for alleles with low protein expression. Even for peptidomes that can be confidently deconvoluted, the pairing between motifs and the expressed MHC alleles is often not trivial, and in many cases must be done manually by visual inspection – with the potential sources of error this process entails.

Here, we have described a fully automated approach, NNAlign_MA, aiming to resolve these challenges. The approach taken in NNAlign_MA is very simple. The method applies a pre-training period where only single allele data (peptide data characterized by having a single MHC association) are included. After this pre-training, the multi-allele data (peptide data characterized by having two or more MHC associations) are annotated using the current prediction model to predict binding to all MHC molecules possible for the peptide, and next defining a single MHC association from the highest prediction value. In this annotation step, multi-allele data are thus casted into a single-allele format, becoming manageable by the NNAlign method and therefore enabled for training. This multi-allele annotation step is iteratively performed in each training cycle.

We have applied the NNAlign_MA method to analyze and interpret three large-scale multi-allele MHC eluted ligand datasets, and demonstrated its unprecedented performance compared to state-of-the-art methods. First, we applied the method to analyze multi-allele HLA MS eluted ligand data from 50 cell lines. Using this data, we demonstrated how the method in the vast majority of cases was capable of correctly identifying distinct binding motifs for each of the HLA molecules expressed in a given cell line. This result is in contrast to findings using earlier methods such as GibbsCluster and MixMHCp that in most cases fail to identify one or more motifs. Also, NNAlign_MA was in close to all cases capable of accurately associating each identified motif with a specific HLA molecule. These results highlighted the high performance of NNAlign_MA compared to current state-of-the-art methods such as MixMHCp/MixMHCpred, where the association of binding motifs to individual HLA molecules is achieved by exclusion principles identifying binding motifs shared uniquely between different cell line datasets.

In terms of predictive performance, the models trained using the NNAlign_MA method were found to outperform conventional methods (trained on single-allele data), both for prediction of HLA eluted ligand data and CD8 epitopes. As expected, this performance gain was most pronounced for ligands/epitopes restricted by HLA molecules characterized by limited or no single-allele data. This observation underlines the single most important power of NNAlign_MA, namely to effectively expand the part of the HLA space covered by accurate predictions. By way of example, the SA EL data included in this study covers 51 HLA molecules. Earlier studies have demonstrated that pan-specific prediction methods allow to accurately predict the binding specificity also for HLA molecules not characterized by binding data, if their distance to a molecule characterized by binding data is 0.1 or less (for a definition of this distance refer to Supplementary Material) (*35*). Applying this rule to the set of 10,558 functional HLA class I A, B and C alleles contained within IPD-IMGT/HLA release 3.35 (*40*) results in a coverage of 76% (8,051 out of 10,558 molecules). By integrating the MA data, the number of alleles covered by EL data is expanded to 85, and number of HLA molecules covered by accurate predictions to 94% (9,949 of 10,558 molecule).

This power of NNAlign_MA to expand the allelic coverage was further demonstrated in a specificity leave-out experiment. Here, entire HLA specificity groups were removed from the single-allele dataset, and the NNAlign_MA framework applied to analyze and characterize multi-allele data including HLA molecules from these removed specificity groups. The result of this experiment confirmed the power of NNAlign_MA to expand the allelic coverage and accurately identifying binding motifs for individual HLA molecules in multi-allele data, also in situations where no explicit information about the binding preferences of the investigated molecules was part of the single-allele training data.

The HLA system has been studied in great detail over the past decades, and peptide-MHC binding data are available for hundreds of alleles. To further explore the predictive power of NNAlign_MA for MHC systems characterized by limited data, we turned to the Bovine Leukocyte Antigen (BoLA) system, and applied NNAlign_MA to analyze MS MHC eluted ligands datasets from 8 cell lines expressing a total of 8 haplotypes covering 16 distinct BoLA molecules. These BoLA molecules shared, for most parts, very low similarity to the molecules included in the single-allele data. Also in this setting, NNAlign_MA was demonstrated to accurately identify binding motifs in all BoLA datasets, and the model trained on the BoLA MA data demonstrated a high predictive power for identification of known BoLA restricted CD8 epitopes, identifying the epitopes within the top 0.3% of the peptides within the epitope source protein sequence. These results thus further demonstrated how NNAlign_MA was capable of correctly deconvoluting binding motifs present in multi-allele data in situations with limited shared similarity to the single-allele data.

As a final validation, the NNAlign_MA framework was applied to MS EL data from MHC II. Also here, the models were evaluated in cross-validation and on an independent set of CD4+ epitopes and the results were in agreement with the results obtained for MHC I. That is, the model trained including MA data showed significantly improved performance compared to models trained on SA data only, when evaluated on both MA EL data and CD4 epitopes.

In a recent work, Bulik-Sullivan et al. (*41*) have suggested an alternative approach to deconvolute and train MHC antigen presentation prediction models using an allele-specific architecture, thus limiting the predictive coverage of the model to MHC alleles present in the training data. This is in contrast to the architecture of NNAlign_MA, which enables pan-specific predictions covering alleles outside the training data (as described above). Also, the allele-specific nature of the method proposed by Bulik-Sullivan et al. limits the power of the tool to identify motifs and construct prediction models for the alleles included in the training data. By way of example in the data presented by Bulik-Sullivan et al., less than 65% of the alleles in their training set ended up covered by a prediction model. Future work and independent benchmarking will allow us to evaluate which of the two approaches is optimal for a given epitope discovery setting.

We have demonstrated how NNAlign_MA achieves binding motif deconvolution driven by similarity to MHC molecules characterized by single-specificity data, and by principles of co-occurrence and exclusion of MHC molecules between different poly-specificity MS eluted ligand dataset. The NNAlign_MA failed in a few cases to construct accurate binding motifs for HLA molecules. These cases were all characterize by very few ligand data, or by alleles only present in single MA data sets. This observation, combined with the power of NNAlign_MA to expand the allelic coverage of the resulting prediction model, points to a direct application NNAlign_MA to effectively achieve broad and high accuracy allelic coverage for regions of the MHC repertoire with yet uncharacterized binding specificities. Guided NNAlign_MA, sets of cell lines with characterized HLA expression should be selected for LC-MS/MS to maximize allele co-occurrence, allele exclusion and allele similarities with the comprehensive set of available EL data so that the NNAlign_MA motif deconvolution for the uncharacterized binding specificities can be achieved in an optimal manner.

While peptide-MHC binding is arguably the most selective step in the MHC antigen presentation pathway, other properties contribute to determining immunogenicity of T cell epitopes. The above mentioned work by Bulik-Sullivan et al. (*41*), attempted to incorporate gene expression levels and proteasome cleavage preferences in a machine-learning model, showing promising improvements for the prediction of cancer neo-epitopes. For the MHC class II system, consistent signatures of peptide trimming and processing have been detected, with pioneering attempts to incorporate them in T cell epitope prediction models (*13, 42*). In future developments of the NNAlign_MA framework, the effect on the predictive power of incorporating such additional potential correlates of immunogenicity will be investigated.

Overall, we have evaluated the proposed NNAlign_MA framework on a large and diverse set of data, and demonstrated how the method in all cases was capable of achieving a complete deconvolution of binding motifs contained within poly-specific MS eluted ligand data, and how the complete deconvolution enabled training prediction models with expanded HLA allelic coverage for accurate identification of both eluted ligands and T cell epitopes. In conclusion, we believe NNAlign_MA offers a universal solution to the challenge of analyzing large-scale MHC peptidomics datasets and consequently affords an optimal way of exploiting the information contained in such data for improving prediction of MHC binding and antigen presentation. The modeling framework is readily extendable to include peptides with post-translational modifications (*15, 43*), and signals from antigen processing located outside the sequence of the ligands (*13*). Given its very high flexibility, we expect NNAlign_MA to serve as an effective tool to further our understanding of the rules for MHC antigen presentation, as a guide for improved T cell epitope discovery and as an aid for effective development of T cell therapeutics.

## Materials and Methods

### Peptide data

Several types of MHC peptide data for human (HLA) and bovine (BoLA) class I, and HLA class II were gathered to train the predictive models presented in this work. Peptide data was classified as single allele data (SA, where each peptide is associated to a single MHC restriction) and multi allele data (MA, where each peptide has multiple options for MHC restriction). MA data are generated from MS MHC ligand elution assays where most often a pan-specific antibody is applied for class I and either a pan-specific class II or a pan-DR specific antibody is applied for class II in the immuno-precipitation step leading to datasets with poly-specificities matching the MHC molecules expressed in the cell line under study. SA data were obtained from binding affinity assays, or from mass spectrometry experiments performed using genetically engineered cell lines that artificially express one single allele.

HLA class I: SA data - both binding affinity (BA), and MS MHC eluted ligands (EL) - was extracted from Jurtz et al. (*14*). The MA data was collected from eight different sources (*12, 25, 26, 28, 44*–*47*). Both datasets were filtered to include only peptides of length 8-14 amino acids. Additional information concerning the HLA class I MA data can be found in Supplementary Table 4 and information concerning the SA BA and EL data sets in Supplementary Table 5.

HLA-II: BA data was extracted from the NetMHCIIpan-3.2 publication (*48*). As for EL data, the Immune Epitope Database *(33*) (IEDB) was queried to identify publications with a large number of allele annotated EL data, both SA and MA (*27, 49*–*56*). Ligands were extracted from these publications, excluding any ligands with post translational modifications. Both BA and EL data was length filtered to include only peptides of length 13-21. Details on the composition of the HLA class II MA data are shown in Supplementary Table 6.

Regarding BoLA, SA data was extracted from Nielsen et al. (*16*) and the MA data was collected either from the same publication (data for the MHC homozygous cell lines expressing the haplotypes A10, A14, A18) or were generated for this study (data for the cell lines expressing the haplotypes A11/A11, A19/A19, A20/A20, A15/A15, and A12/A15). The novel EL data were generated as described earlier (*16*) by use of LC-MS using the pan anti-BoLA class I antibody IL-88 for immunoprecipitation. Sequence interpretation of MS/MS spectra was performed using a database containing all bovine protein entries in Uniprot combined (in cases of *Theileria parva* infected cell lines) with 4084 annotated *Theileria parva* protein sequences (*57*). Spectral interpretation was performed using PEAKS 7.5 (*58*) (Bioinformatics Solutions). All datasets were filtered to include only peptides of length 8-14. A summary of the BoLA MA data is given in Supplementary Table 7.

### Training data

Three training sets were constructed, one for each of the systems under study (Table 2). To ensure an unbiased performance evaluation on the MA data, duplicated entries between the SA EL and MA data were first removed from the SA EL dataset for each training set. Next, random peptides were extracted from the UniProt database and used as negative instances for the EL data in each case. Here, an equal amount of random negatives was used for each length, consisting of five times the amount of peptides for the most abundant length in the given positive EL dataset as described earlier (*13, 16*). This enrichment with random natural negative peptides was done for each individual SA and MA EL dataset. The amount of positive and negative peptides in each training set is shown in Table 2.

**Table 2.**
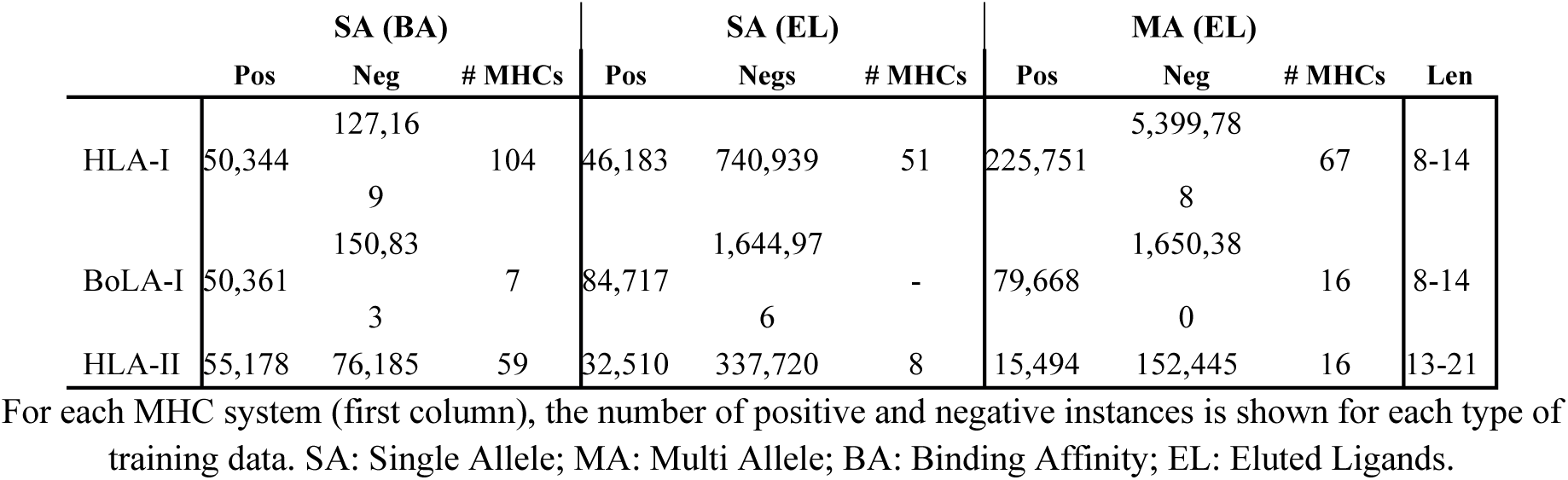
Training data overview.

### Evaluation data

For HLA class I, an independent evaluation data set of HLA restricted CD8+ epitopes was obtained from Jurtz et al. (*14*). After removal of epitopes overlapping with the HLA-I training data, the final evaluation data consisted of 558 HLA-epitope entries.

For the evaluation with MixMHCpred, MHCFlurry, and MHCFlurry_EL (the version of MHCFLurry trained including EL data), the epitope data set was further filtered to only include epitopes restricted to HLA molecules covered by all method. This resulted in a data set of 541 epitopes. Since MixMHCpred cannot make predictions for peptides containing X, all such peptides were removed from the benchmark prior to evaluation.

For BoLA-I, a set of BoLA restricted epitopes were obtained from Nielsen et al. (*16*). For HLA-I and BoLA-I evaluation, the source protein sequence of each epitope was *in-silico* digested into overlapping 8-14mers, and the performance reported as the Frank score, i.e. proportion of peptides with a prediction score higher than that of the epitope (*14*). Using this measure, a value of 0 corresponds to a perfect prediction (the known epitope is identified with the highest predicted binding value among all peptides found within the source protein) and a value of 0.5 to random prediction.

For HLA class II all CD4+ epitopes measured by Intracellular Cytokine Staining(ICS) assay were downloaded from the IEDB. The set was filtered to include only positive epitopes with four letter resolution HLA typing. Furthermore, epitopes overlapping with the HLA-II training data (100% identity) were removed. As for HLA-I, the Frank was used to validate the model performance, here in-silico digesting the source protein into overlapping of a length equal to that of the epitope. Finally, to exclude potential noise, epitopes were discarded if none of the predictions methods included in the benchmark could identify the epitope with a Frank value of 0.2 or less. This resulted in a set of 221 HLA-II epitopes for evaluation.

### NNAlign modeling and training hyperparameters

Models for peptide-MHC binding prediction were trained with hyperparameters and model architectures similar to those described earlier (*13, 15, 29*) for prediction of peptide-MHC binding based on the datasets described in Table 2. Positive instances in EL datasets (for both SA and MA) were labeled with a target value of 1, and negatives with a target value of 0.

In order to avoid performance overestimation and model overfitting, training sets were split into 5 partitions for cross-validation purposes using the common motif algorithm (*59*) with a motif length of 8 amino acids for class I (corresponding to the shortest binding mode for class I peptides) and 9 amino acids for class II (corresponding to the size of the class II binding core) as described earlier (*13, 14*).

A single and simple yet highly critical step sets the updated NNAlign_MA method proposed here aside from its ancestors. To be able to accurately handle and annotate MA data, NNAlign_MA imposes a burn-in period where the method is trained only on SA data. After the burn-in period, each data point in the MA dataset is annotated by predicting binding to all possible MHC molecules defined in the MA dataset, and assigning the restriction from the highest prediction value (see prediction score rescaling for exceptions from this setup). After this annotation step, the SA and MA data are merged respecting the data partitioning to further train the algorithm.

In the case of HLA-I and BoLA-I, models were trained on the full set of SA and MA data as an ensemble of 50 individual networks, generated from 5 different seeds; 56 and 66 hidden neurons; and 5 partitions for cross-validation. Models were trained for 200 iterations (using early stopping), with a burn-in period of 20 iterations. For performance comparison, a SA-only model was trained for HLA-I using the same architecture and hyper-parameters by excluding all MA data from the cross-validation partitions, thus including only data for SA while respecting the training data structure.

Regarding HLA-II, default settings for MHC-II prediction as previously described (*13, 29*) were used. Models were trained and evaluated on 5-fold cross-validation partitions defined by common motif clustering with a motif of length 9. The final ensemble of models consists of 250 networks (2, 10, 20, 40 and 60 hidden neurons and 10 random weight initiation seeds for each CV fold). Networks were trained for 400 iterations, without early stopping and using a burn-in period of 20.

### Prediction score rescaling

To level out differences in the prediction scores between MHC alleles imposed by the differences in number of positive training examples and distance to the training data included in the SA dataset, a rescaling of the raw prediction values was implemented and applied in the MA data annotation. The rescaling was implemented as a z-score transformation of the raw prediction values using the relation 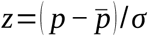, where *p* is the raw prediction value of the peptide to a given MHC molecules, and 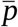 and *σ* are the mean and standard deviation of the distribution of prediction values for random natural peptides for the MHC molecule. Here, the score distribution was estimated by predicting binding of 10,000 random natural 9mer peptides to MHC molecule in question. Next, the mean and standard deviation were estimated from the right-hand side of the distribution, iteratively excluding outliers (z-score < −3 or z-score > 3). This estimation of 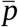 and *σ* was repeated in each iteration round prior to annotating the MA data. As the rescaling is imposed to level out score differences between MHC molecules characterized in the SA training binding data and molecules from the MA data distant to the training data, the need for rescaling should be leveled out as the MA data are included in the training and the NNAlign_MA training progresses. To achieve this, the values of 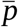 and *σ* were modified to converge towards uniform values 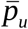 and *σ* _*u*_ defined as the average of 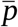 and *σ* over all molecules in the MA dataset. This convergence was defined as 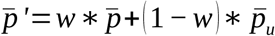 and 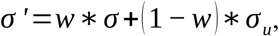 where *w*=1/(1+*e*^(*x*−75) /10^) and *x* is the number of training iterations. With this relation, *w* is close to 1 after pre-training (x = 20), and converges to 0 as x passes 100 iterations.

### Distance between pairs of MHC molecules

The distance, D, between MHC molecules was calculated as described earlier (*32*) from the sequence similarity between the pseudo sequences of the two molecules. Likewise, was the distance of an MHC molecule to the data used to train a given prediction model, defined as the closest distance to any MHC molecule included in the training data.

### Pruning the HLA supertype tree - HLA models with removed specificities

In order to quantify how MA data can boost the performance of a peptide-MHC predictor, we constructed additional models, where SA data associated with HLA molecules from the A2 and A3 supertypes where exclude from the training data. In short, this was achieved by first identifying the alleles in the MA data for the two supertypes (*36*), resulting in the following allele list: HLA-A*02:01, HLA-A*02:05, HLA-A*02:06, HLA-A*02:20, HLA-A*68:02, HLA-A*03:01, HLA-A*11:01, HLA-A*30:01, HLA-A*31:01, and HLA-A*68:01. Next, all data for alleles with a distance (see above) of less than 0.2 to any of the alleles in this list were removed from the SA data. Finally, an SA model was trained as described above on the remaining SA data, and MA model on the remaining SA data combined with the complete MA data, respecting the original data partitioning.

### Sequence motif construction

Sequence binding motif were visualized as Kullback-Leibler logo plots using Seg2Logo (*60*). Amino acids are grouped by negatively charged (red), positively charged (blue), polar (green) or hydrophobic (black). If not otherwise specified, binding motifs were generated from the top 0.1% of 200,000 random natural peptides (9mers for class I and 15mers for class II) as described earlier^14^.

### Binding motifs similarity comparison

The similarity between two HLA binding motifs was estimated in terms of the Pearson’s correlation coefficient (PCC) between the two vectors of 9*20 elements (9 positions and 20 amino acid propensity scores at each position).

### Model performance evaluation

For model comparison, the AUC (Area Under the ROC Curve) and AUC 0.1 (Area Under the ROC Curve integrated up to a False Positive Rate of 10%) performance measures were used. For a given model, each test set was predicted using the model trained during cross-validation. Next, all test sets were concatenated, and an AUC/AUC0.1 value was calculated for each MHC molecule/cell line identifier. In case of multi-allele data, the prediction score to each peptide was assigned as the maximal prediction value over the set of possible MHC molecules.

To evaluate the “cleanness” of a given cluster/motif identified by NNAlign_MA, positive-predictive values (PPV) were calculated. For each cell line, we calculated the number of ligands N predicted to be bound to each allele from the concatenated test set predictions. Next, the PPV for each motif was calculated as the fraction of peptides in the top N*0.95 predictions that were actual ligands. The values of 95% was selected to tolerate a certain proportion of noise in the EL data (*29*).

### *In vitro* binding data

Recombinnt BoLA-1*00901 and human beta-2 microglobulins (β2m) were produced as previously described (*61*). In brief, biotinylated BoLA-1*00901 was generated in Escherichia coli, harvested as inclusion bodies, extracted into Tris-buffered 8 M urea and purified using ion exchange, hydrophobic, and gel filtration chromatographies. MHC-I heavy chain proteins were never exposed to reducing conditions, which allows for purification of highly active pre-oxidized BoLA molecules, which folds efficiently when diluted into an appropriate reaction buffer. The pre-oxidized, denatured proteins were stored at −20 °C in Tris-buffered 8 M urea. Human β2m was expressed and purified as previously described (*62*).

Nonameric peptide binding motifs were determined for BoLA-1*00901, using PSCPL as previously described (*34, 61, 63*). Recombinant, biotinylated BoLA heavy chain molecules in 8 M urea were diluted at least 100-fold into PBS buffer containing 125I-labeled human β2m and peptide to initiate pMHC-I complex formation. The final concentration of BoLA was between 10 and 100 nM, depending on the specific activity of the heavy chain. The reactions were carried out in the wells of streptavidin-coated scintillation 384-well FlashPlate® PLUS microplates (Perkin Elmer, Waltham, MA). Recombinant radiolabeled human β2m and saturating concentrations (10 μM) of peptide were allowed to reach steady state by overnight incubation at 18 °C. After overnight incubation, excess unlabeled bovine β2m was added to a final concentration of 1 μM and the temperature was raised to 37 °C to initiate dissociation. pMHC-I dissociation was monitored for 24 h by consecutive measurement of the scintillation microplate on a scintillation TopCount NXT multiplate counter (Perkin Elmer, Waltham, MA). PSCPL dissociation data were analyzed as described^33^. Briefly, following background correction, the area under the dissociation curve (AUC) was calculated for each sublibrary by summing the counts from 0 to 24 h. The relative contribution of each residue in each position (i.e., the relative binding, RB) was calculated as RB=(AUC_sublibrary/AUC_X9). The RB values were next normalized to sum to one for each peptide position and used as input to Seq2Logo to generate the *in vitro* BoLA-1*00901 binding-motif.

### The BoLA benchmark

The NNAlign_MA framework was used to train a model on the SA data combined with MA data from the total of 8 BoLA cell lines (for details on these datasets, see above). After training, we proceeded to investigate the different binding motifs captured by the model. In particular, we were interested in the motifs of the BoLA molecules shared between multiple haplotypes. One such example is BoLA-2*02501, present in the A14, and A15 haplotypes. While the motif for this molecule in our earlier work showed a strong preference for G/Q and L at P2 and P9 respectively (Figure 7B, the new NNAlign framework captured a completely different signal, with a consistent proline (P) signal on most positions (see Figure 7A). Also, the number of ligands assigned to BoLA-2*02501 was extremely low for the three cell lines expressing this molecule (less than 1.5% in all three cases). A prime source of this result is the very large distance (0.426) of BoLA-2*02501 to the SA training data (for details on this distance measure refer to section “Distance between pairs of MHC molecules” above). In comparison, the maximum distance between any molecule in the MA data to the SA training data for the HLA system is less than 0.13. These large pairwise distances in the BoLA system have two strong impacts on the predictive behavior of NNAlign_MA. First and foremost, the model pre-trained on the SA is expected to have limited power to predict the binding motif of the BoLA-2*02501 molecule (in the pre-trained model, BoLA-2*02501 has a preference for P at P2). Secondly, the prediction values of the pre-trained model will be lower for this molecule compared to molecules that share higher similarity to the SA data used for pre-training. To deal with the latter of these two issues, we devised a rescaling scheme for the prediction score in the MA annotation step of the NNAlign_MA framework, and rescaled the raw prediction by comparing it to a score distribution obtained from a large set of random natural peptides (for details section “Prediction score rescaling” above). This score distribution was recalculated prior to each MA annotation. Including this rescaling step, the number of ligands estimated by cross-validation to be assigned to BoLA-2*02501 from the three cell lines increased to 13% on average. However, investigating the motifs from the ligands predicted to be associated with BoLA-2*02501 from the A14 MA data to that from A12/A15 and A15 MA data, an inconsistency became apparent (see Figure 7C and D). Here, the motif obtained from the A14 MA data showed an additional preference for G at P2 that was completely absent for the motif obtained from the A15 and A12/A15 MA data.

**Figure 7.**
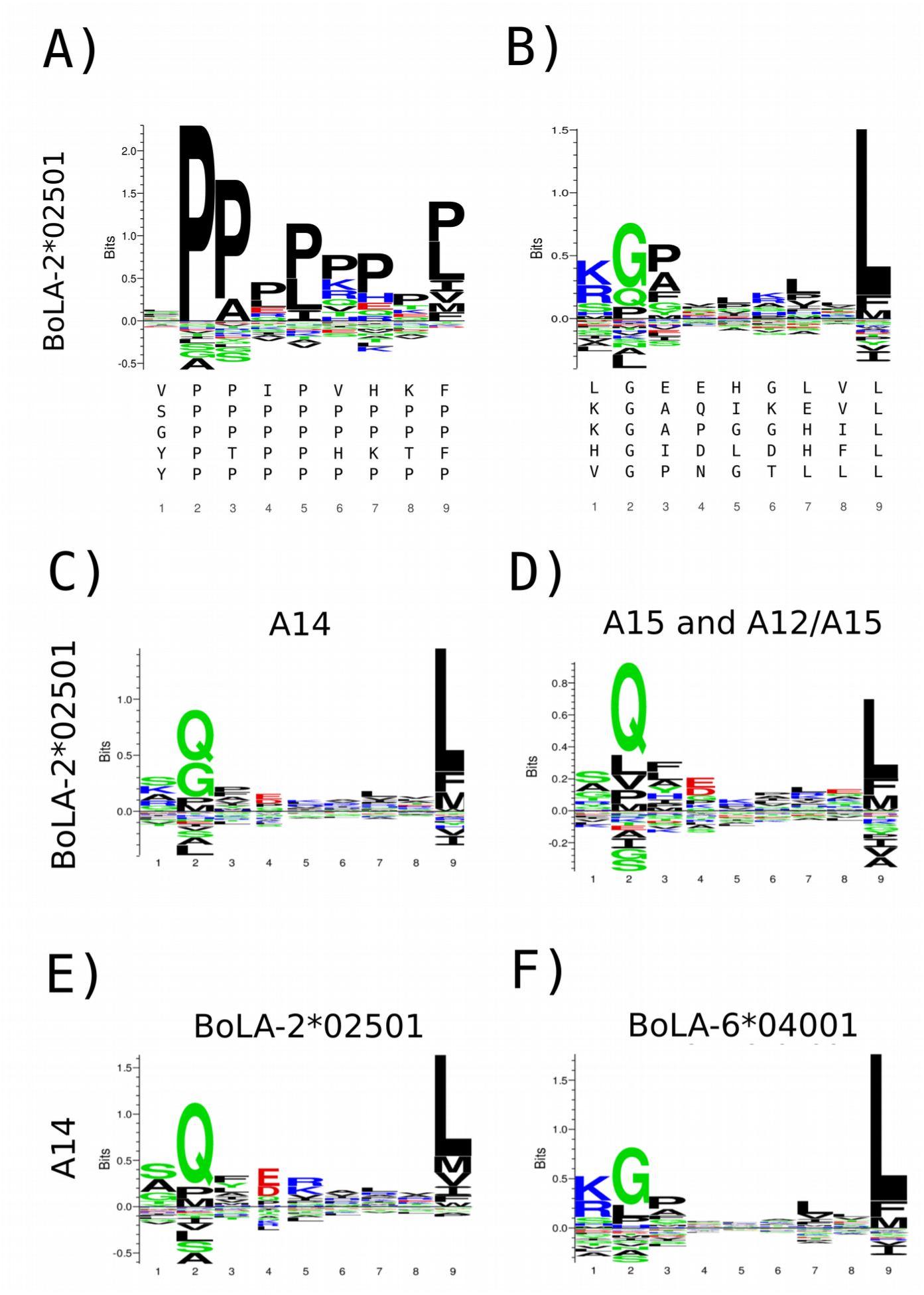
Figure 7. Identification of binding motifs for the BoLA-2*02501 molecule A) Binding preference for this molecule found by the NNAlign_MA method without score rescaling, B) motif logo found in our previous work (*16*). The top five repeating binding cores present in each motif alignment are shown below each logo. Binding motifs for BoLA-2*02501 obtained by NNAlign_MA trained including three BoLA alleles (BoLA-1*02301, BoLA-4*02401, and BoLA-2*02501) for the A14 MA data from ligands predicted using cross-validation to be restricted by BoLA-2*02501 from the A14 MA data (C), and from the A15 and A12/15 MA datasets (D). Binding motifs for the BoLA-2*02501 (E) and BoLA-6*04001 (F) molecules as estimated from ligands in the A14 MA data as predicted by NNAlign_MA using cross-validation when expanding the list of alleles in A14 to include BoLA-6*04001.

Re-examining the original publication that described the BoLA allele expression profile in the A14 haplotype (*64*), suggested an explanation to the apparent inconsistencies in the predicted BoLA-2*02501 binding motifs. In that paper, A14 was found to express 4 and not 3 BoLA alleles, as was assumed in our earlier publication and used in the first NNAlign_MA analysis. The extra allele expressed is BoLA-6*04001. After including this extra allele in the A14 haplotype and retraining the model, we obtained the binding motifs displayed in Figure 1E, showing a motif for BoLA-2*02501 consistent with the motif identified in the A12/A15 and A15 MA data (Supplementary Material Figure 7D), and a likewise well-defined motif for BoLA-6*04001 (Figure 1F). These results clearly suggest that the motif earlier reported for BoLA-2*02501 (Figure 7B) was a mixture of the motifs of BoLA-2*02501 and BoLA-6*04001.

## Acknowledgments

This work was supported in part by the Federal funds from the National Institute of Allergy and Infectious Diseases, National Institutes of Health, Department of Health and Human Services, under Contract No. HHSN272201200010C, Bill and Melinda Gates Foundation [OPP1078791]; and by the Science and Technology Council of Investigation (CONICET-Argentina).

## Author contributions

MN designed the study. BA, BR, and MN performed the experiments and statistical analysis. SB, NT and TC generated the experimental data. BA, BR, CB, MA and MN analyzed and interpreted the data and wrote the paper. All authors read and approved the final manuscript.

