## Supplementary Data for "NNAlign_MA; MHC peptidome deconvolution for accurate MHC binding motif characterization and improved T cell epitope predictions"

**Supplementary Figure 1.** Full NNAlign_MA motif deconvolution for the Multi Allele (MA) HLA-I data analyzed in this work.

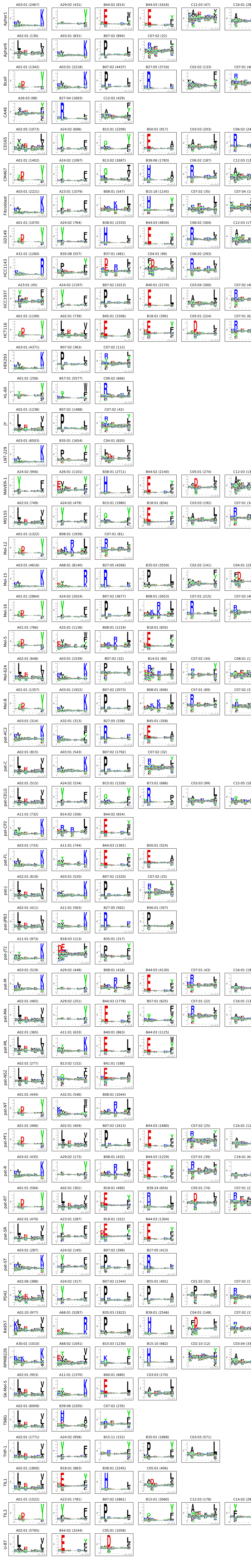

Each row corresponds to a cell line present in the training data (50 in total; for more details, refer to Supplementary Table 4). Using cross validation, each ligand is assigned to one of the HLA alleles expressed in the given cell line. Using this assignment, binding motifs were generated for each allele in each cell line using Seq2Logo (60). To remove potential MS contaminants, only ligands with a prediction score greater than 0.01 were included. Above each logo is given the number of sequences associated to the corresponding HLA allele. For details on the accuracy of this clustering, refer to Supplementary Figures 2 and 3.

**Supplementary Figure 2.** Comparison between NNAlign_MA deconvoluted motifs and motifs derived from single-allele (SA) data for alleles in the multi-allele data set where SA data are available from the IEDB (33).

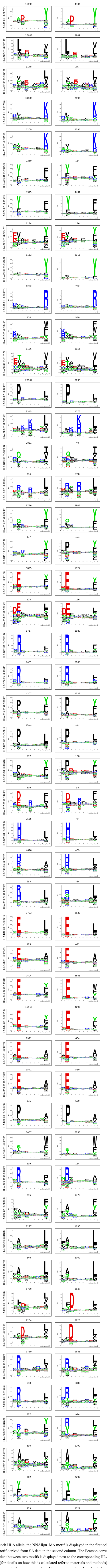

For each HLA allele, the NNAlign_MA motif is displayed in the first column; the motif derived from SA data in the second column. The Pearson correlation coefficient between two motifs is displayed next to the corresponding HLA name (for details on how this is calculated refer to materials and methods). The amount of sequences employed to construct a given logo is displayed on top of each logo.

**Supplementary Figure 3.** Correlation matrices between NNAlign_MA motifs for HLA alleles that are shared between five or more cell lines.

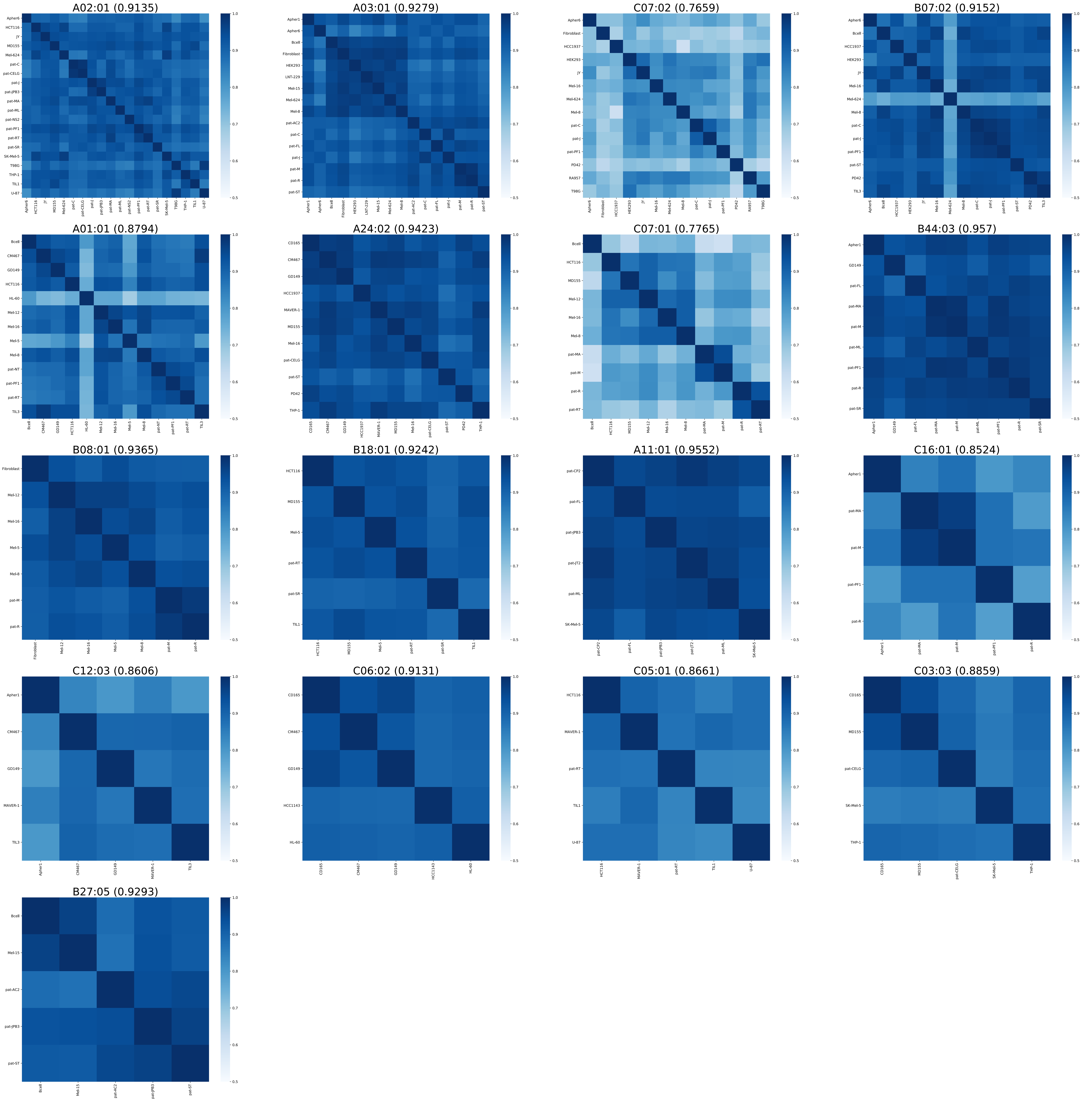


Each matrix displays the Pearson correlation coefficients between all motifs found by NNAlign_MA for a given HLA allele, across all the cell lines sharing the allele (for details on how the correlation is calculated refer to materials and methods). The average Pearson correlation coefficient for each matrix is given next to the corresponding allele name.

**Supplementary Figure 4.** Full NNAlign_MA motif deconvolution for the Multi Allele (MA) BoLA-I data analyzed in this work.

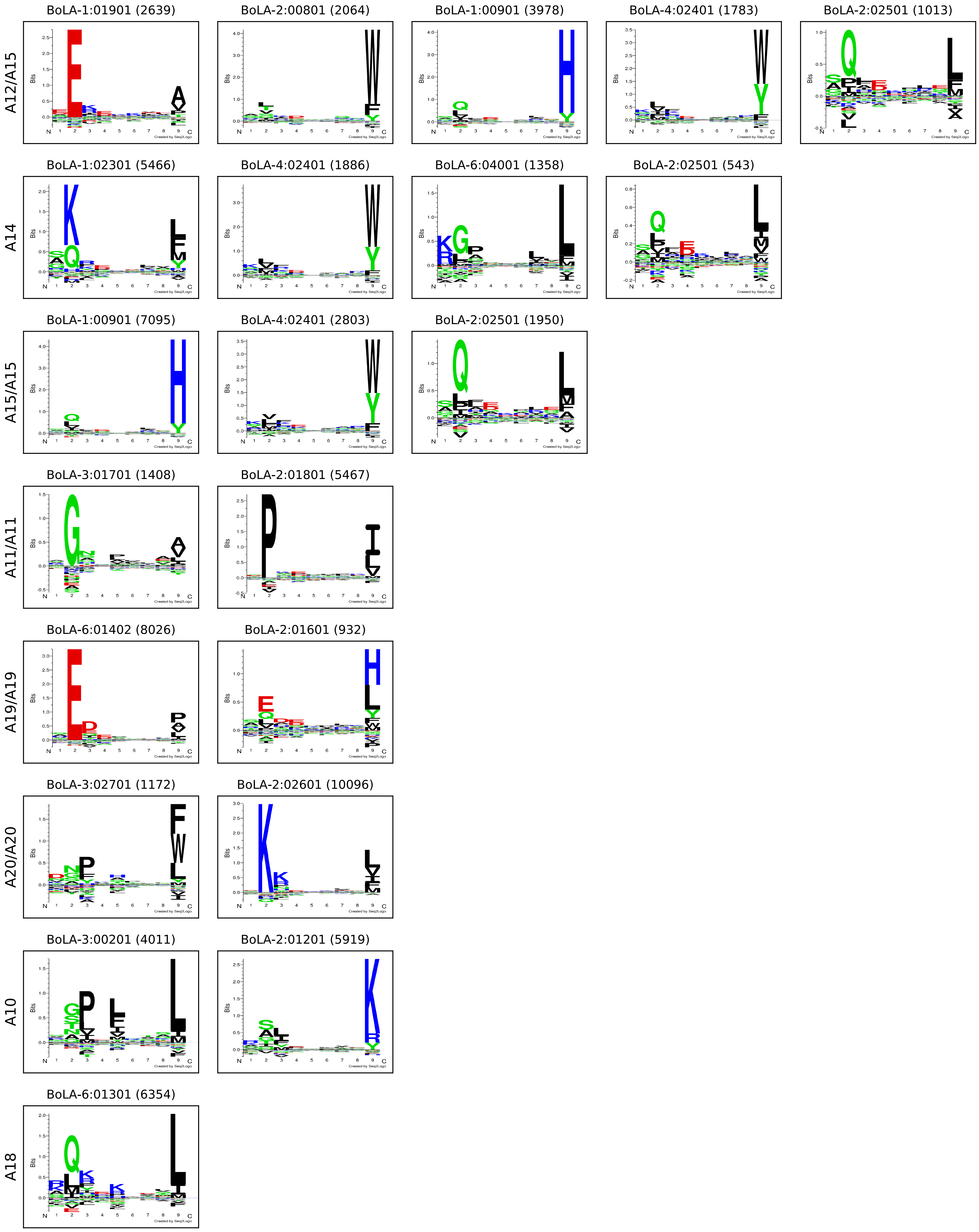


Each row corresponds to a MA data set present in the training data. Using cross validation, each ligand is assigned to one of the BoLA alleles expressed in the given data set. Using this assignment, binding motifs were generated for each allele in each cell line using Seq2Logo (60). To remove potential MS contaminants, only ligands with a prediction score greater than 0.01 were included. Above each logo is given the number of sequences associated to the corresponding BoLA allele.

**Supplementary Figure 5.** Full NNAlign_MA motif deconvolution for the Multi Allele (MA) HLA-II data analyzed in this work.

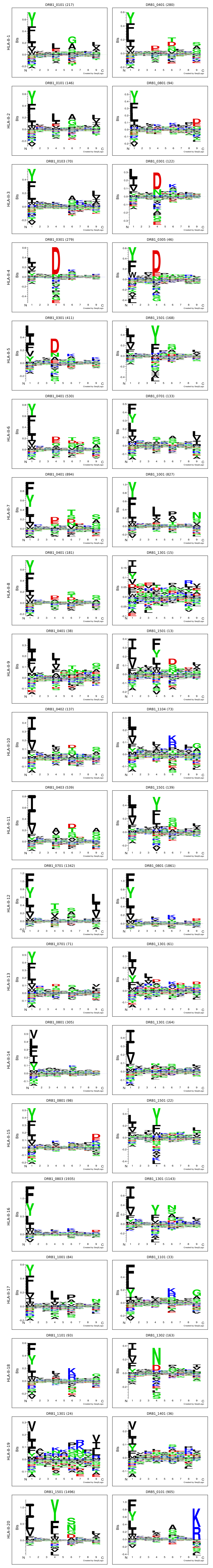


Each row corresponds to a MA data set present in the training data. Using cross validation, each ligand is assigned to one of the HLA alleles expressed in the given data set. Using this assignment, binding motifs were generated for each allele in each cell line using Seq2Logo (60). To remove potential MS contaminants, only ligands with a prediction score greater than 0.01 were included. Above each logo is given the number of sequences associated to the corresponding HLA allele.

**Supplementary Table 1.** Predicted Positive Predictive Values (PPV) for all clusters associated with each allele in each cell line the multi allele (MA) dataset after NNAlign_MA deconvolution. For details on the calculation of PPV, refer to materials and methods.

| Cell Line ID | HLA Allele | PPV |
|---|---|---|
| Apher1 | HLA-A03:01 | 0,84 |
| | HLA-A29:02 | 0,73 |
| | HLA-B44:02 | 0,81 |
| | HLA-B44:03 | 0,91 |
| | HLA-C12:03 | 0,75 |
| | HLA-C16:01 | 0,49 |
| Apher6 | HLA-A02:01 | 0,74 |
| | HLA-A03:01 | 0,82 |
| | HLA-B07:02 | 0,91 |
| | HLA-C07:02 | 0,38 |
| Bcell | HLA-A01:01 | 0,79 |
| | HLA-A03:01 | 0,75 |
| | HLA-B07:02 | 0,88 |
| | HLA-B27:05 | 0,83 |
| | HLA-C02:02 | 0,53 |
| | HLA-C07:01 | 0,39 |
| CA46 | HLA-A26:03 | 0,35 |
| | HLA-B27:04 | 0,90 |
| | HLA-C12:02 | 0,77 |
| CD165 | HLA-A02:05 | 0,64 |
| | HLA-A24:02 | 0,81 |
| | HLA-B15:01 | 0,87 |
| | HLA-B50:01 | 0,82 |
| | HLA-C03:03 | 0,75 |
| | HLA-C06:02 | 0,83 |
| CM467 | HLA-A01:01 | 0,90 |
| | HLA-A24:02 | 0,84 |
| | HLA-B13:02 | 0,80 |
| | HLA-B39:06 | 0,75 |
| | HLA-C06:02 | 0,83 |
| | HLA-C12:03 | 0,69 |
| Fibroblast | HLA-A03:01 | 0,84 |
| | HLA-A23:01 | 0,84 |
| | HLA-B08:01 | 0,67 |
| | HLA-B15:18 | 0,88 |
| | HLA-C07:02 | 0,61 |
| | HLA-C07:04 | 0,50 |
| GD149 | HLA-A01:01 | 0,79 |
| | HLA-A24:02 | 0,78 |
| | HLA-B38:01 | 0,90 |
| | HLA-B44:03 | 0,92 |
| | HLA-C06:02 | 0,82 |
| | HLA-C12:03 | 0,54 |
| HCC1143 | HLA-A31:01 | 0,84 |
| | HLA-B35:08 | 0,82 |
| | HLA-B37:01 | 0,77 |

| | | |
|---|---|---|
| | HLA-C04:01 | 0,59 |
| | HLA-C06:02 | 0,80 |
| HCC1937 | HLA-A23:01 | 0,55 |
| | HLA-A24:02 | 0,87 |
| | HLA-B07:02 | 0,83 |
| | HLA-B40:01 | 0,90 |
| | HLA-C03:04 | 0,78 |
| | HLA-C07:02 | 0,35 |
| HCT116 | HLA-A01:01 | 0,86 |
| | HLA-A02:01 | 0,74 |
| | HLA-B18:01 | 0,86 |
| | HLA-B45:01 | 0,89 |
| | HLA-C05:01 | 0,71 |
| | HLA-C07:01 | 0,59 |
| HEK293 | HLA-A03:01 | 0,90 |
| | HLA-B07:02 | 0,71 |
| | HLA-C07:02 | 0,54 |
| HL-60 | HLA-A01:01 | 0,38 |
| | HLA-B57:01 | 0,92 |
| | HLA-C06:02 | 0,81 |
| JY | HLA-A02:01 | 0,85 |
| | HLA-B07:02 | 0,91 |
| | HLA-C07:02 | 0,48 |
| LNT-229 | HLA-A03:01 | 0,82 |
| | HLA-B35:01 | 0,69 |
| | HLA-C04:01 | 0,47 |
| MAVER-1 | HLA-A24:02 | 0,85 |
| | HLA-A26:01 | 0,81 |
| | HLA-B38:01 | 0,92 |
| | HLA-B44:02 | 0,90 |
| | HLA-C05:01 | 0,54 |
| | HLA-C12:03 | 0,62 |
| MD155 | HLA-A02:01 | 0,76 |
| | HLA-A24:02 | 0,84 |
| | HLA-B15:01 | 0,88 |
| | HLA-B18:01 | 0,87 |
| | HLA-C03:03 | 0,72 |
| | HLA-C07:01 | 0,45 |
| Mel-12 | HLA-A01:01 | 0,77 |
| | HLA-B08:01 | 0,91 |
| | HLA-C07:01 | 0,63 |
| Mel-15 | HLA-A03:01 | 0,76 |
| | HLA-A68:01 | 0,81 |
| | HLA-B27:05 | 0,76 |
| | HLA-B35:03 | 0,82 |
| | HLA-C02:02 | 0,48 |
| | HLA-C04:01 | 0,38 |
| Mel-16 | HLA-A01:01 | 0,77 |
| | HLA-A24:02 | 0,85 |
| | HLA-B07:02 | 0,89 |
| | HLA-B08:01 | 0,87 |
| | HLA-C07:01 | 0,80 |
| | HLA-C07:02 | 0,52 |

| | | |
|---|---|---|
| Mel-5 | HLA-A01:01 | 0,52 |
| | HLA-A25:01 | 0,82 |
| | HLA-B08:01 | 0,82 |
| | HLA-B18:01 | 0,87 |
| Mel-624 | HLA-A02:01 | 0,82 |
| | HLA-A03:01 | 0,86 |
| | HLA-B07:02 | 0,31 |
| | HLA-B14:01 | 0,71 |
| | HLA-C07:02 | 0,62 |
| | HLA-C08:01 | 0,25 |
| Mel-8 | HLA-A01:01 | 0,84 |
| | HLA-A03:01 | 0,79 |
| | HLA-B07:02 | 0,90 |
| | HLA-B08:01 | 0,84 |
| | HLA-C07:01 | 0,75 |
| | HLA-C07:02 | 0,54 |
| PD42 | HLA-A02:06 | 0,58 |
| | HLA-A24:02 | 0,88 |
| | HLA-B07:02 | 0,91 |
| | HLA-B55:01 | 0,82 |
| | HLA-C01:02 | 0,53 |
| | HLA-C07:02 | 0,48 |
| RA957 | HLA-A02:20 | 0,64 |
| | HLA-A68:01 | 0,83 |
| | HLA-B35:03 | 0,87 |
| | HLA-B39:01 | 0,89 |
| | HLA-C04:01 | 0,72 |
| | HLA-C07:02 | 0,53 |
| RPMI8226 | HLA-A30:01 | 0,66 |
| | HLA-A68:02 | 0,79 |
| | HLA-B15:03 | 0,77 |
| | HLA-B15:10 | 0,91 |
| | HLA-C02:10 | 0,60 |
| | HLA-C03:04 | 0,73 |
| SK-Mel-5 | HLA-A02:01 | 0,81 |
| | HLA-A11:01 | 0,85 |
| | HLA-B40:01 | 0,86 |
| | HLA-C03:03 | 0,57 |
| T98G | HLA-A02:01 | 0,78 |
| | HLA-B39:06 | 0,65 |
| | HLA-C07:02 | 0,43 |
| THP-1 | HLA-A02:01 | 0,81 |
| | HLA-A24:02 | 0,83 |
| | HLA-B15:11 | 0,63 |
| | HLA-B35:01 | 0,84 |
| | HLA-C03:03 | 0,76 |
| TIL1 | HLA-A02:01 | 0,78 |
| | HLA-B18:01 | 0,81 |
| | HLA-B38:01 | 0,91 |
| | HLA-C05:01 | 0,53 |
| TIL3 | HLA-A01:01 | 0,86 |
| | HLA-A23:01 | 0,80 |
| | HLA-B07:02 | 0,86 |
| | HLA-B15:01 | 0,85 |

| | | |
|---|---|---|
| | HLA-C12:03 | 0,61 |
| | HLA-C14:02 | 0,68 |
| U-87 | HLA-A02:01 | 0,78 |
| | HLA-B44:02 | 0,81 |
| | HLA-C05:01 | 0,65 |
| pat-AC2 | HLA-A03:01 | 0,79 |
| | HLA-A32:01 | 0,79 |
| | HLA-B27:05 | 0,78 |
| | HLA-B45:01 | 0,84 |
| pat-C | HLA-A02:01 | 0,51 |
| | HLA-A03:01 | 0,58 |
| | HLA-B07:02 | 0,78 |
| | HLA-C07:02 | 0,65 |
| pat-CELG | HLA-A02:01 | 0,60 |
| | HLA-A24:02 | 0,82 |
| | HLA-B15:01 | 0,80 |
| | HLA-B73:01 | 0,70 |
| | HLA-C03:03 | 0,61 |
| | HLA-C15:05 | 0,61 |
| pat-CP2 | HLA-A11:01 | 0,87 |
| | HLA-B14:02 | 0,80 |
| | HLA-B44:02 | 0,93 |
| pat-FL | HLA-A03:01 | 0,77 |
| | HLA-A11:01 | 0,83 |
| | HLA-B44:03 | 0,89 |
| | HLA-B50:01 | 0,78 |
| pat-J | HLA-A02:01 | 0,66 |
| | HLA-A03:01 | 0,70 |
| | HLA-B07:02 | 0,79 |
| | HLA-C07:02 | 0,51 |
| pat-JPB3 | HLA-A02:01 | 0,85 |
| | HLA-A11:01 | 0,82 |
| | HLA-B27:05 | 0,83 |
| | HLA-B56:01 | 0,72 |
| pat-JT2 | HLA-A11:01 | 0,89 |
| | HLA-B18:03 | 0,74 |
| | HLA-B35:01 | 0,78 |
| pat-M | HLA-A03:01 | 0,63 |
| | HLA-A29:02 | 0,61 |
| | HLA-B08:01 | 0,68 |
| | HLA-B44:03 | 0,86 |
| | HLA-C07:01 | 0,44 |
| | HLA-C16:01 | 0,59 |
| pat-MA | HLA-A02:01 | 0,72 |
| | HLA-A29:02 | 0,64 |
| | HLA-B44:03 | 0,92 |
| | HLA-B57:01 | 0,69 |
| | HLA-C07:01 | 0,53 |
| | HLA-C16:01 | 0,62 |
| pat-ML | HLA-A02:01 | 0,70 |
| | HLA-A11:01 | 0,77 |
| | HLA-B40:01 | 0,89 |
| | HLA-B44:03 | 0,93 |

| | | |
|---|---|---|
| pat-NS2 | HLA-A02:01 | 0,87 |
| | HLA-B13:02 | 0,74 |
| | HLA-B41:01 | 0,71 |
| pat-NT | HLA-A01:01 | 0,74 |
| | HLA-A32:01 | 0,75 |
| pat-PF1 | HLA-A01:01 | 0,71 |
| | HLA-A02:01 | 0,72 |
| | HLA-B07:02 | 0,89 |
| | HLA-B44:03 | 0,92 |
| | HLA-C07:02 | 0,33 |
| | HLA-C16:01 | 0,45 |
| pat-R | HLA-A03:01 | 0,57 |
| | HLA-A29:02 | 0,61 |
| | HLA-B08:01 | 0,65 |
| | HLA-B44:03 | 0,86 |
| | HLA-C07:01 | 0,83 |
| | HLA-C16:01 | 0,48 |
| pat-RT | HLA-A01:01 | 0,69 |
| | HLA-A02:01 | 0,71 |
| | HLA-B18:01 | 0,85 |
| | HLA-B39:24 | 0,79 |
| | HLA-C05:01 | 0,54 |
| | HLA-C07:01 | 0,44 |
| pat-SR | HLA-A02:01 | 0,73 |
| | HLA-A23:01 | 0,80 |
| | HLA-B18:01 | 0,85 |
| | HLA-B44:03 | 0,91 |
| pat-ST | HLA-A03:01 | 0,82 |
| | HLA-A24:02 | 0,86 |
| | HLA-B07:02 | 0,89 |
| | HLA-B27:05 | 0,84 |

**Supplementary Table 2.** Cell lines with atypical HLA-A or HLA-B peptidome repertoire profiles.

| Cell line ID | HLA-A | HLA-B | HLA-C | Source PMID(s) |
|---|---|---|---|---|
| HL-60 | 0.047 | 0.851 | 0.102 | 26992070<br>17083564 |
| CA46 | 0.071 | 0.739 | 0.191 | 26992070 |
| Mel-624 | 0.931 | 0.048 | 0.021 | 7541714 |
| HEK293 | 0.896 | 0.080 | 0.024 | 26258424 |

For each cell line, the relative peptidome size for the HLA-A, HLA-B and HLA-C loci is given. Peptidome sizes were calculated as described in Figure 3 B. The last column shows the references to earlier publications describing the loss of the locus for a given cell line.

**Supplementary Table 3.** AUC0.1 performance values for the SA molecules left out from the training of the A2 and A3 specificity reduced SA and MA models.

| | SA MODEL | MA MODEL |
|---|---|---|
| HLA-A*02:01 | 0.376 | 0.778 |
| HLA-A*02:05 | 0.533 | 0.771 |
| HLA-A*02:07 | 0.477 | 0.871 |
| HLA-A*03:01 | 0.359 | 0.896 |
| HLA-A*11:01 | 0.361 | 0.862 |
| HLA-A*26:01 | 0.838 | 0.962 |
| HLA-A*29:01 | 0.762 | 0.873 |
| HLA-A*31:01 | 0.409 | 0.749 |
| HLA-A*32:01 | 0.808 | 0.905 |

Note that not all the molecules included in the evaluation are part of the A2 and A3 supertypes; these molecules are included because they have a distance to the A2 and A3 molecules in the MA dataset less than 0.1.

**Supplementary Table 4.** Summary of the multi-allele (MA) data included in the HLA-I benchmark.

| Cell line ID | Positives | Negatives | HLA-A | | HLA-B | | HLA-C | | Source PMID |
|---|---|---|---|---|---|---|---|---|---|
| CD165 | 5364 | 132869 | HLA-A*02:05 | HLA-A*24:02 | HLA-B*15:01 | HLA-B*50:01 | HLA-C*03:03 | HLA-C*06:02 | 28832583 |
| CM467 | 7401 | 184646 | HLA-A*01:01 | HLA-A*24:02 | HLA-B*13:02 | HLA-B*39:06 | HLA-C*06:02 | HLA-C*12:03 | |
| GD149 | 9756 | 208444 | HLA-A*01:01 | HLA-A*24:02 | HLA-B*38:01 | HLA-B*44:03 | HLA-C*06:02 | HLA-C*12:03 | |
| MD155 | 4374 | 108036 | HLA-A*02:01 | HLA-A*24:02 | HLA-B*15:01 | HLA-B*18:01 | HLA-C*03:03 | HLA-C*07:01 | |
| PD42 | 2577 | 48693 | HLA-A*02:06 | HLA-A*24:02 | HLA-B*07:02 | HLA-B*55:01 | HLA-C*01:02 | HLA-C*07:02 | |
| RA957 | 11037 | 232658 | HLA-A*02:20 | HLA-A*68:01 | HLA-B*35:03 | HLA-B*39:01 | HLA-C*04:01 | HLA-C*07:02 | |
| TIL1 | 5445 | 140312 | HLA-A*02:01 | HLA-A*02:01 | HLA-B*18:01 | HLA-B*38:01 | HLA-C*05:01 | - | |
| TIL3 | 8799 | 206212 | HLA-A*01:01 | HLA-A*23:01 | HLA-B*07:02 | HLA-B*15:01 | HLA-C*12:03 | HLA-C*14:02 | |
| Apher1 | 6145 | 123349 | HLA-A*03:01 | HLA-A*29:0 | HLA-B*44:02 | HLA-B*44:03 | HLA-C*12:03 | HLA-C*16:01 | |
| Apher6 | 1962 | 39798 | HLA-A*02:01 | HLA-A*03:01 | HLA-B*07:02 | - | HLA-C*07:02 | - | |
| Mel-15 | 21813 | 395324 | HLA-A*03:01 | HLA-A*68:01 | HLA-B*27:05 | HLA-B*35:03 | HLA-C*02:02 | HLA-C*04:01 | 27869121 |
| Mel-16 | 11980 | 264233 | HLA-A*01:01 | HLA-A*24:02 | HLA-B*07:02 | HLA-B*08:01 | HLA-C*07:01 | HLA-C*07:02 | |
| Mel-12 | 3758 | 88425 | HLA-A*01:01 | HLA-A*01:01 | HLA-B*08:01 | - | HLA-C*07:01 | - | |
| Mel-8 | 6251 | 119000 | HLA-A*01:01 | HLA-A*03:01 | HLA-B*07:02 | HLA-B*08:01 | HLA-C*07:01 | HLA-C*07:02 | |
| Mel-5 | 4749 | 106896 | HLA-A*01:01 | HLA-A*25:01 | HLA-B*08:01 | HLA-B*18:01 | - | - | |
| Fibroblast | 5289 | 122127 | HLA-A*03:01 | HLA-A*23:01 | HLA-B*08:01 | HLA-B*15:18 | HLA-C*07:02 | HLA-C*07:04 | 25576301 |
| HCC1143 | 2780 | 69565 | HLA-A*31:01 | - | HLA-B*35:08 | HLA-B*37:01 | HLA-C*04:01 | HLA-C*06:02 | |
| HCC1937 | 4976 | 102331 | HLA-A*23:01 | HLA-A*24:02 | HLA-B*07:02 | HLA-B*40:01 | HLA-C*03:04 | HLA-C*07:02 | |
| HCT116 | 4174 | 93208 | HLA-A*01:01 | HLA-A*02:01 | HLA-B*45:01 | HLA-B*18:01 | HLA-C*05:01 | HLA-C*07:01 | |
| JY | 2868 | 60863 | HLA-A*02:01 | - | HLA-B*07:02 | - | HLA-C*07:02 | - | |
| Bcell | 12199 | 220971 | HLA-A*01:01 | HLA-A*03:01 | HLA-B*07:02 | HLA-B*27:05 | HLA-C*02:02 | HLA-C*07:01 | 24616531 |
| Mel-624 | 2375 | 49050 | HLA-A*02:01 | HLA-A*03:01 | HLA-B*07:02 | HLA-B*14:01 | HLA-C*07:02 | HLA-C*08:01 | 27600516 |
| SK-Mel-5 | 3293 | 64537 | HLA-A*02:01 | HLA-A*11:01 | HLA-B*40:01 | - | HLA-C*03:03 | - | |
| HEK293 | 4972 | 86634 | HLA-A*03:01 | - | HLA-B*07:02 | - | HLA-C*07:02 | - | 27600516 |
| MAVER-1 | 7403 | 171783 | HLA-A*24:02 | HLA-A*26:01 | HLA-B*38:01 | HLA-B*44:02 | HLA-C*05:01 | HLA-C*12:03 | |
| HL-60 | 6607 | 115694 | HLA-A*01:01 | - | HLA-B*57:01 | - | HLA-C*06:02 | - | |
| RPMI8226 | 4524 | 113201 | HLA-A*30:01 | HLA-A*68:02 | HLA-B*15:03 | HLA-B*15:10 | HLA-C*02:10 | HLA-C*03:04 | |
| THP-1 | 5542 | 142866 | HLA-A*02:01 | HLA-A*24:02 | HLA-B*15:11 | HLA-B*35:01 | HLA-C*03:03 | - | |
| CA46 | 2324 | 62647 | HLA-A*26:03 | - | HLA-B*27:04 | - | HLA-C*12:02 | - | |
| LNT-229 | 10311 | 177908 | HLA-A*03:01 | - | HLA-B*35:01 | - | HLA-C*04:01 | - | 27412690 |
| T98G | 10011 | 216072 | HLA-A*02:01 | - | HLA-B*39:06 | - | HLA-C*07:02 | - | |
| U-87 | 11585 | 241396 | HLA-A*02:01 | - | HLA-B*44:02 | - | HLA-C*05:01 | - | |
| pat-AC2 | 1369 | 32168 | HLA-A*03:01 | HLA-A*32:01 | HLA-B*27:05 | HLA-B*45:01 | - | - | 27841757 |
| pat-C | 2983 | 49759 | HLA-A*02:01 | HLA-A*03:01 | HLA-B*07:02 | - | HLA-C*07:02 | - | |
| pat-CELG | 3814 | 72328 | HLA-A*02:01 | HLA-A*24:02 | HLA-B*15:01 | HLA-B*73:01 | HLA-C*03:03 | HLA-C*15:05 | |
| pat-CP2 | 1790 | 36895 | HLA-A*11:01 | - | HLA-B*14:02 | HLA-B*44:02 | - | - | |
| pat-FL | 3629 | 74392 | HLA-A*03:01 | HLA-A*11:01 | HLA-B*44:03 | HLA-B*50:01 | - | - | |
| pat-J | 2552 | 42497 | HLA-A*02:01 | HLA-A*03:01 | HLA-B*07:02 | - | HLA-C*07:02 | - | |
| pat-JPB3 | 1937 | 35295 | HLA-A*02:01 | HLA-A*11:01 | HLA-B*27:05 | HLA-B*56:01 | - | - | |
| pat-JT2 | 1467 | 29587 | HLA-A*11:01 | - | HLA-B*18:03 | HLA-B*35:01 | - | - | |
| pat-M | 2476 | 53262 | HLA-A*03:01 | HLA-A*29:02 | HLA-B*08:01 | HLA-B*44:03 | HLA-C*07:01 | HLA-C*16:01 | |
| pat-MA | 3682 | 69891 | HLA-A*02:01 | HLA-A*29:02 | HLA-B*44:03 | HLA-B*57:01 | HLA-C*07:01 | HLA-C*16:01 | |
| pat-ML | 3139 | 55262 | HLA-A*02:01 | HLA-A*11:01 | HLA-B*40:01 | HLA-B*44:03 | - | - | |
| pat-NS2 | 636 | 15212 | HLA-A*02:01 | - | HLA-B*13:02 | HLA-B*41:01 | - | - | |
| pat-NT | 2190 | 53238 | HLA-A*01:01 | HLA-A*32:01 | HLA-B*08:01 | - | - | - | |
| pat-PF1 | 4646 | 86859 | HLA-A*01:01 | HLA-A*02:01 | HLA-B*07:02 | HLA-B*44:03 | HLA-C*07:02 | HLA-C*16:01 | |
| pat-R | 2372 | 49169 | HLA-A*03:01 | HLA-A*29:02 | HLA-B*08:01 | HLA-B*44:03 | HLA-C*07:01 | HLA-C*16:01 | |
| pat-RT | 2537 | 49846 | HLA-A*01:01 | HLA-A*02:01 | HLA-B*18:01 | HLA-B*39:24 | HLA-C*05:01 | HLA-C*07:01 | |
| pat-SR | 2632 | 57417 | HLA-A*02:01 | HLA-A*23:01 | HLA-B*18:01 | HLA-B*44:03 | - | - | |
| pat-ST | 1256 | 26963 | HLA-A*03:01 | HLA-A*24:02 | HLA-B*07:02 | HLA-B*27:05 | - | - | |

“Positives” and “Negatives” refer to the number of positive and negative instances contained in each cell line data. Further rows show the HLA-A, HLA-B and HLA-C expressed by a given cell line, together with the Source ID for its corresponding dataset.

**Supplementary Table 5.** Single Allele (SA) Binding Affinity (BA) and Eluted Ligands (EL) training data summary for the HLA-I system. For the BA training set, the total amount of sequences per MHC molecule (discarding artificial negatives) is shown; in the case of EL data, the total amount of positives is displayed.

| BA data | |
|---|---|
| **Alleles** | **# Peptides** |
| HLA-A01:01 | 3985 |
| HLA-A02:01 | 11097 |
| HLA-A02:02 | 3613 |
| HLA-A02:03 | 5728 |
| HLA-A02:04 | 3 |
| HLA-A02:05 | 66 |
| HLA-A02:06 | 4802 |
| HLA-A02:07 | 66 |
| HLA-A02:10 | 18 |
| HLA-A02:11 | 1081 |
| HLA-A02:12 | 1181 |
| HLA-A02:16 | 919 |
| HLA-A02:17 | 341 |
| HLA-A02:19 | 1243 |
| HLA-A02:50 | 134 |
| HLA-A03:01 | 6513 |
| HLA-A03:02 | 26 |
| HLA-A03:19 | 30 |
| HLA-A11:01 | 5614 |
| HLA-A11:02 | 14 |
| HLA-A23:01 | 1871 |
| HLA-A24:02 | 2315 |
| HLA-A24:03 | 1374 |
| HLA-A25:01 | 959 |
| HLA-A26:01 | 3729 |
| HLA-A26:02 | 641 |
| HLA-A26:03 | 535 |
| HLA-A29:02 | 2286 |
| HLA-A30:01 | 2711 |
| HLA-A30:02 | 1564 |
| HLA-A31:01 | 5152 |
| HLA-A32:01 | 908 |
| HLA-A32:07 | 88 |
| HLA-A32:15 | 74 |
| HLA-A33:01 | 2508 |
| HLA-A66:01 | 207 |
| HLA-A68:01 | 3190 |
| HLA-A68:02 | 4542 |
| HLA-A68:23 | 82 |
| HLA-A69:01 | 2559 |
| HLA-A74:01 | 15 |
| HLA-A80:01 | 1167 |
| HLA-B07:02 | 4100 |
| HLA-B08:01 | 3171 |

| EL data | |
|---|---|
| **Alleles** | **# Peptides** |
| HLA-A01:01 | 3405 |
| HLA-A02:01 | 5349 |
| HLA-A02:05 | 98 |
| HLA-A02:07 | 30 |
| HLA-A03:01 | 427 |
| HLA-A11:01 | 313 |
| HLA-A24:02 | 2699 |
| HLA-A24:06 | 169 |
| HLA-A24:13 | 49 |
| HLA-A26:01 | 97 |
| HLA-A29:02 | 4377 |
| HLA-A31:01 | 31 |
| HLA-A32:01 | 29 |
| HLA-B07:02 | 3324 |
| HLA-B08:01 | 458 |
| HLA-B13:01 | 57 |
| HLA-B15:01 | 455 |
| HLA-B15:02 | 52 |
| HLA-B18:01 | 46 |
| HLA-B27:02 | 2334 |
| HLA-B27:03 | 278 |
| HLA-B27:04 | 569 |
| HLA-B27:05 | 2567 |
| HLA-B27:06 | 646 |
| HLA-B27:07 | 1253 |
| HLA-B27:08 | 1306 |
| HLA-B27:09 | 1363 |
| HLA-B35:01 | 680 |
| HLA-B35:03 | 23 |
| HLA-B35:08 | 93 |
| HLA-B37:01 | 39 |
| HLA-B39:06 | 495 |
| HLA-B40:01 | 1286 |
| HLA-B40:02 | 1548 |
| HLA-B41:01 | 19 |
| HLA-B41:03 | 55 |
| HLA-B41:04 | 37 |
| HLA-B44:02 | 1662 |
| HLA-B44:03 | 303 |
| HLA-B44:27 | 24 |
| HLA-B44:28 | 18 |
| HLA-B45:01 | 150 |
| HLA-B49:01 | 119 |
| HLA-B50:01 | 114 |

| | |
|---|---|
| HLA-B08:02 | 1018 |
| HLA-B08:03 | 469 |
| HLA-B14:01 | 42 |
| HLA-B14:02 | 283 |
| HLA-B15:01 | 4213 |
| HLA-B15:02 | 164 |
| HLA-B15:03 | 604 |
| HLA-B15:09 | 830 |
| HLA-B15:17 | 1444 |
| HLA-B15:42 | 364 |
| HLA-B18:01 | 2370 |
| HLA-B27:01 | 4 |
| HLA-B27:02 | 8 |
| HLA-B27:03 | 874 |
| HLA-B27:04 | 4 |
| HLA-B27:05 | 3372 |
| HLA-B27:06 | 7 |
| HLA-B27:10 | 2 |
| HLA-B27:20 | 92 |
| HLA-B35:01 | 2724 |
| HLA-B35:03 | 93 |
| HLA-B35:08 | 1 |
| HLA-B37:01 | 50 |
| HLA-B38:01 | 500 |
| HLA-B39:01 | 1785 |
| HLA-B40:01 | 2984 |
| HLA-B40:02 | 712 |
| HLA-B40:13 | 59 |
| HLA-B42:01 | 160 |
| HLA-B42:02 | 18 |
| HLA-B44:02 | 1954 |
| HLA-B44:03 | 1006 |
| HLA-B45:01 | 627 |
| HLA-B45:06 | 362 |
| HLA-B46:01 | 1798 |
| HLA-B48:01 | 881 |
| HLA-B51:01 | 2383 |
| HLA-B52:01 | 12 |
| HLA-B53:01 | 1341 |
| HLA-B54:01 | 731 |
| HLA-B57:01 | 2640 |
| HLA-B57:02 | 18 |
| HLA-B57:03 | 34 |
| HLA-B58:01 | 3116 |
| HLA-B58:02 | 56 |
| HLA-B73:01 | 122 |
| HLA-B81:01 | 26 |
| HLA-B83:01 | 339 |
| HLA-C03:03 | 153 |
| HLA-C04:01 | 552 |
| HLA-C05:01 | 172 |
| HLA-C06:02 | 309 |
| HLA-C07:01 | 241 |
| HLA-C07:02 | 142 |

| | |
|---|---|
| HLA-B51:01 | 2424 |
| HLA-B57:01 | 370 |
| HLA-C03:04 | 29 |
| HLA-C04:01 | 366 |
| HLA-C05:01 | 435 |
| HLA-C07:02 | 19 |
| HLA-C16:01 | 222 |
| H2-Db | 808 |
| H2-Kb | 1906 |
| H2-Kd | 663 |
| Mamu-B008:01 | 495 |

| | |
|---|---|
| HLA-C08:02 | 87 |
| HLA-C12:03 | 172 |
| HLA-C14:02 | 259 |
| HLA-C15:02 | 252 |
| HLA-E01:01 | 96 |
| HLA-E01:03 | 55 |
| Mamu-A01 | 2466 |
| Mamu-A02 | 1188 |
| Mamu-A07 | 535 |
| Mamu-A11 | 1144 |
| Mamu-A20102 | 132 |
| Mamu-A2201 | 582 |
| Mamu-A2601 | 142 |
| Mamu-A70103 | 95 |
| Mamu-B01 | 444 |
| Mamu-B03 | 973 |
| Mamu-B04 | 2 |
| Mamu-B08 | 952 |
| Mamu-B1001 | 140 |
| Mamu-B17 | 1384 |
| Mamu-B3901 | 439 |
| Mamu-B52 | 846 |
| Mamu-B6601 | 101 |
| Mamu-B8301 | 368 |
| Mamu-B8701 | 144 |
| Patr-A0101 | 337 |
| Patr-A0301 | 262 |
| Patr-A0401 | 233 |
| Patr-A0602 | 1 |
| Patr-A0701 | 495 |
| Patr-A0901 | 621 |
| Patr-B0101 | 636 |
| Patr-B0901 | 1 |
| Patr-B1301 | 196 |
| Patr-B1701 | 8 |
| Patr-B2401 | 293 |
| SLA-10401 | 185 |
| SLA-10701 | 23 |
| SLA-20401 | 105 |
| SLA-30401 | 76 |
| BoLA-AW10 | 166 |
| BoLA-D18.4 | 258 |
| BoLA-HD6 | 268 |
| BoLA-JSP.1 | 158 |
| BoLA-T2a | 167 |
| BoLA-T2b | 157 |
| BoLA-T2C | 90 |
| Gogo-B0101 | 14 |
| H-2-Db | 2580 |
| H-2-Dd | 276 |
| H-2-Kb | 3694 |
| H-2-Kd | 811 |
| H-2-Kk | 364 |
| H-2-Ld | 260 |

| | |
|---|---|
| H-2-Lq | 2 |

**Supplementary Table 6.** Multi Allele (MA) data summary for the HLA-II benchmark.

| ID | Positives | Negatives | HLA-DRB | | Source PMID |
|---|---|---|---|---|---|
| HLA-II-1 | 509 | 5355 | DRB1*01:01 | DRB1*04:01 | 27726376 |
| HLA-II-2 | 240 | 2655 | DRB1*01:01 | DRB1*08:01 | 27726376 |
| HLA-II-3 | 200 | 1890 | DRB1*01:03 | DRB1*03:01 | 27726376 |
| HLA-II-4 | 327 | 3285 | DRB1*03:01 | DRB1*03:05 | 27726376 |
| HLA-II-5 | 595 | 6930 | DRB1*03:01 | DRB1*15:01 | 27452731, 27726376 |
| HLA-II-6 | 670 | 7740 | DRB1*04:01 | DRB1*07:01 | 27452731 |
| HLA-II-7 | 1772 | 17460 | DRB1*04:01 | DRB1*10:01 | 27726376 |
| HLA-II-8 | 213 | 2520 | DRB1*04:01 | DRB1*13:01 | 27452731 |
| HLA-II-9 | 51 | 480 | DRB1*04:01 | DRB1*15:01 | 27726376 |
| HLA-II-10 | 210 | 2475 | DRB1*04:02 | DRB1*11:04 | 27726376 |
| HLA-II-11 | 682 | 7335 | DRB1*04:03 | DRB1*15:01 | 27726376 |
| HLA-II-12 | 3216 | 29565 | DRB1*07:01 | DRB1*08:01 | 29632711 |
| HLA-II-13 | 145 | 1710 | DRB1*07:01 | DRB1*13:01 | 27452731 |
| HLA-II-14 | 496 | 4860 | DRB1*08:01 | DRB1*13:01 | 27452731, 29632711 |
| HLA-II-15 | 121 | 1440 | DRB1*08:01 | DRB1*15:01 | 27726376 |
| HLA-II-16 | 3080 | 29745 | DRB1*08:03 | DRB1*13:01 | 29632711 |
| HLA-II-17 | 118 | 1215 | DRB1*10:01 | DRB1*11:01 | 27726376 |
| HLA-II-18 | 257 | 2835 | DRB1*11:01 | DRB1*13:02 | 27726376 |
| HLA-II-19 | 65 | 585 | DRB1*13:01 | DRB1*14:01 | 27452731 |
| HLA-II-20 | 2426 | 22365 | DRB1*15:01 | DRB5*01:01 | 28467828 |

"Positives" and "Negatives" refer to the number of positive and negative instances contained in each MA EL data set. Further rows show the HLA-DRB alleles expressed by a given cell line, together with the Source PMID(s) for its corresponding dataset.

**Supplementary Table 7.** Multi Allele (MA) data summary for the BoLA benchmark.

<table>
<tr><th>Cell line ID</th><th>Haplotype</th><th>Positives</th><th>Negatives</th><th>BoLA-1</th><th>BoLA-2</th><th>BoLA-3</th><th>BoLA-4</th><th>BoLA-6</th><th>Source PMID</th></tr>
<tr><td rowspan="2">2123</td><td rowspan="2">A12/A15</td><td rowspan="2">11908</td><td rowspan="2">310336</td><td>BoLA-1*01901</td><td>BoLA-2*00801</td><td rowspan="2">-</td><td rowspan="2">BoLA-4*02401</td><td rowspan="2">-</td><td rowspan="6">-</td></tr>
<tr><td>BoLA-1*00901</td><td>BoLA-2*02501</td></tr>
<tr><td>5072</td><td>A11</td><td>7469</td><td>153959</td><td>-</td><td>BoLA-2*01801</td><td>BoLA-3*01701</td><td>-</td><td>-</td></tr>
<tr><td>2824</td><td>A19</td><td>9665</td><td>175580</td><td>-</td><td>BoLA-2*01601</td><td>-</td><td>-</td><td>BoLA-6*01402</td></tr>
<tr><td>5350</td><td>A20</td><td>11791</td><td>274531</td><td>-</td><td>BoLA-2*02601</td><td>BoLA-3*02701</td><td>-</td><td>-</td></tr>
<tr><td>2408</td><td>A15</td><td>12523</td><td>320919</td><td>BoLA-1*00901</td><td>BoLA-2*02501</td><td></td><td>BoLA-4*02401</td><td>-</td></tr>
<tr><td>1011/500004</td><td>A10</td><td>10188</td><td>148801</td><td>-</td><td>BoLA-2*01201</td><td>BoLA-3*00201</td><td>-</td><td>-</td><td rowspan="3">29115832</td></tr>
<tr><td>641</td><td>A18</td><td>6615</td><td>80170</td><td>-</td><td>-</td><td>-</td><td>-</td><td>BoLA-6*01301</td></tr>
<tr><td>2229/104003</td><td>A14</td><td>9509</td><td>186084</td><td>BoLA-1*02301</td><td>BoLA-2*02501</td><td>-</td><td>BoLA-4*02401</td><td>BoLA-6*04001</td></tr>
</table>

“Positives” and “Negatives” refer to the number of positive and negative instances contained in each cell line data. Further rows show the BoLA-1, BoLA-2, BoLA-3, BoLA-4, and BoLA-6 alleles expressed by a given cell line, together with the Source PMID for its corresponding dataset. Allele annotation was obtained from Vasoya, D. et al. (*64*)
